# Hyper-specialized primates possess a reduced suite of xenobiotic-metabolizing cytochrome P450 genes

**DOI:** 10.1101/2023.12.06.570463

**Authors:** Morgan E. Chaney, Anthony J. Tosi, Christina M. Bergey

## Abstract

Subfamilies of cytochrome P450 proteins have been strongly linked to the metabolism of physiologically disruptive compounds such as alkaloids, terpenoids, and other xenobiotics. Consistent with this function, these genes have adaptively evolved in response to environmental pressures exerted on animals, such as herbivores, that consume elevated amounts of toxic xenobiotics or plant secondary metabolites (PSMs). Theory on evolutionary tradeoffs predicts that highly specialized herbivores should exhibit a relatively narrow toolkit of adaptations to accommodate the concomitantly narrow arrays of PSMs in their diets. The bamboo lemurs of Madagascar (genera *Prolemur* and *Hapalemur*) represent an interesting test case for this theory because of their dietary hyper-specialization, as these lemurs consume bamboo and grasses at rates otherwise unseen in the order Primates. To test whether the hyper-specialized folivory of these primates is reflected in a similarly specialized and narrow P450 gene suite, we assembled a dataset of confidently assembled *CYP1-3* genes for two species of bamboo lemur as well as additional lemur species. We tested the predictions that bamboo lemurs would exhibit, first, greater rates of gene loss for xenobiotic-metabolizing P450s and, second, relaxed selection on xenobiotic-metabolizing P450 subfamilies relative to lemurs without such dietary hyper-specialization. We found support for the first prediction, related to gene loss, in the *CYP2B, CYP2C, CYP2D, CYP2J,* and *CYP3A* subfamilies, all of which encode xenobiotic metabolizers. We additionally inferred relaxation of selection for the *CYP2F* and *CYP2J* subfamilies. The evolution of the P450 genes in bamboo lemurs provides support for the evolutionary tradeoff hypothesis, and we further hypothesize that, rather than adapting to a general array of PSMs, bamboo lemurs have instead adapted to the primary toxin in their diet, the highly potent poison cyanide.

**Highlights:**

- Some of the most specialized diets among primates are those of bamboo lemurs.
- Bamboo lemurs have fewer xenobiotic-metabolizing P450 genes than other lemurs.
- Natural selection has relaxed on the *CYP2F* and *CYP2J* subfamilies in bamboo lemurs.

## 1. Introduction

Among mammals, a large proportion of xenobiotic compounds are transformed or detoxified by enzymes within the cytochrome P450 superfamily. Because of this role that P450s play in mediating potential environmental toxins and because species vary in their diets and exposure to such compounds, selective pressure has induced non-random additive and adaptive changes in genetic sequence and copy number among species (Chaney et al., 2018, 2020; Ingelman-Sundberg, 2022; Kawashima & Satta, 2014; Kirischian et al., 2011; Skopec et al., 2022). The effect of evolving toxin resistance has been especially well-documented for pest insect species because of the economic importance of pesticides and insects’ evolutionary responses to those substances (Wondji et al. 2009; Feyereisen 2011; Bass et al. 2014; Wang et al. 2014; Walsh et al. 2018; Joussen and Heckel 2021). Studies of P450 gene adaptation outside of pest insects have often focused on dietary xenobiotics or plant secondary metabolites (PSMs). For example, research on a model system comprising several species of woodrat (*Neotoma* spp.) has shown that PSMs common in their diet (*e.g.*, alkaloids, terpenoids) are metabolized especially by CYP2B enzymes and, to a minor extent, other xenobiotic-metabolizing enzymes (Haley et al., 2007a, 2007b; Orr et al., 2020; Wilderman et al., 2014).

Comparisons between specialist and generalist species of these woodrats have shed light on the effects that diet can have on P450 evolution. For instance, the generalist woodrat species (*N. albigula*) was found to have lower copy number but greater sequence diversity of *CYP2B* relative to a specialist (*N. stephensi*). This greater diversity of the generalist is consistent with the broader array of PSMs with which the generalist must contend (Kitanovic et al., 2018). The increased copy number of the specialist, in contrast, is in line with the prediction that genes important for coping with the specialist’s narrow diet will be subject to *additive* evolutionary processes (*e.g.*, gene duplication, positive selection) that will enhance the species’ ability to mitigate a narrow range of metabolically harmful compounds (Dearing et al., 2005; Freeland & Janzen, 1974). This is taken to an extreme for specialized koala (*Phascolartcos cinereus*), in which the key xenobiotic metabolizing *CYP2C* subfamily has reached a copy number of 31 via serial duplication (Johnson et al., 2018).

Mammalian dietary specialization has also been linked to patterns of gene loss (Blumer et al., 2022; Hecker et al., 2019; Itoigawa et al., 2019; Lauterbur, 2019; Marciniak et al., 2021; Wagner et al., 2022). For example, vampire bats (genera *Desmodus, Diaemus,* and *Diphylla*) have lost sweet and umami taste receptors (Zhao et al., 2010b; Zhao et al., 2012) and have experienced reduction of their bitter-taste sense through pseudogenization of *TAS2R* genes (Hong & Zhao, 2014). Interestingly, losses of gustatory genes have also been found for cetaceans, which have lost nearly all taste-receptor genes apart from those for salty taste (Feng et al., 2014), and for pandas (*Ailuropoda melanoleuca*), which have lost their umami taste-receptors (Zhao et al., 2010a). Theory predicts that genes unimportant to a specialist’s dietary adaptation will be subject to a suite of *subtractive* evolutionary processes (*e.g.,* accumulation of deleterious mutations, pseudogenization, or loss of protein coding region or entire gene). Gene loss could either be for adaptive reasons (see Olson, 1999) or because of a release of selective pressure (see Blumer et al., 2022; Huelsmann et al., 2019). Despite these examples and the theory surrounding them, subtractive mechanisms such as gene loss have remained relatively understudied compared to additive mechanisms such as gene duplication and positive or directional selection.

In summary, specialized folivory is predicted to have two evolutionary effects on a species’ xenobiotic-metabolizing toolkit: First, genes involved in detoxification or avoidance of xenobiotic toxins that are rich in a folivorous specialist’s diet should show signals of selection in the form of positive copy-number changes or positive selection. Second, genes *not* involved in such species’ defense against toxins in their specialized diet should exhibit signals of negative copy-number changes and relaxation of selection. Although the first prediction is well-supported by prior studies of P450 gene evolution, comparatively few studies have investigated patterns of P450 gene loss or relaxation of selection in specialized folivores.

Bamboo lemurs represent a well-suited case for testing predictions about P450 gene loss in specialized folivores due to their narrow diets, which are among the most specialized in the entire primate order. Despite their diminutive body sizes (0.9-2.6 kg: Tan, 2006), they inhabit an ecological niche that is often compared with that of the giant panda (Eronen et al., 2017; McKenney et al., 2017; Tan, 2006) and are the only extant primates known to specialize so intensely on bamboo or grass species (Table S1). The three sympatric species at Ranomafana National Park (*i.e*., *Hapalemur aureus, H. griseus*, and *Prolemur simus*) are the source of much of the current understanding of bamboo lemur ecology. All three of these populations have diets that are often predominantly composed of a small number of species of bamboo or liana (Grassi, 2006; Overdorff et al., 1997; Tan, 1999), and the sympatry of these species has been understood in terms of niche partitioning, notably by their differential reliance on separate plant parts from the same species of giant bamboo (Glander et al., 1989). Although other species outside of Ranomafana (*e.g.*, *H. meridionalis, H. alaotrensis*) do not actually consume bamboo, they nonetheless eat grasses (Poaceae) or sedges (Cyperaceae) at similarly high frequencies (Table S1). A recent report, also based on locations outside of Ranomafana, demonstrated more dietary flexibility for *Prol. simus* than would have previously been expected (Mihaminekena et al., 2024; see Table S1), but bamboo is nonetheless critical forthis species’ evolution, likely as a staple fallback food (Marshall & Wrangham, 2007), as evidenced by alimentary traits that are likely related to a specialized folivorous (dentition: Jernvall et al., 2008; Vinyard et al., 2008; gastrointestinal morphology: Yamashita et al., 2008; M.E. Chaney, pers. obs.; see Campbell et al., 2000). Regardless, all three populations subsisted on diets composed of 90% folivorous foods or more, even if one population’s diet was only 41.5% bamboo (Mihaminekena et al., 2024). Thus, regardless of habitat or species, bamboo lemurs subsist on highly folivorous diets that are notably low in their diversity.

One bamboo PSM is particularly intriguing for its toxicity. Consumption of the specific bamboo parts consumed by *H. aureus* and *Prol. simus* exposes these lemurs to high levels of potent cyanogenic glycosides (Ballhorn et al., 2009; Eppley et al., 2017; Glander et al., 1989). When plant tissue is disrupted by some destructive process like mastication, these glycosides will be hydrolyzed by the plant cell’s own ꞵ-glycosidase enzymes to release cyanide (Ballhorn, 2011; Cressey & Reeve, 2019). At Ranomafana, two bamboo lemur species have been estimated to consume many times the estimated lethal dose of cyanide for mammals of their body sizes (Ballhorn et al., 2009; Glander et al., 1989). Although cyanide, the primary known antifeedant of this bamboo, is detoxified by a metabolic pathway that does not include any cytochrome P450 enzymes (reviewed by Isom et al., 2015), bamboo lemurs likely have to contend with other PSMs known to be present in bamboo species, such as phenolic and polyphenolic compounds (Zhu et al. 2018). Thus, we still expect evolution of bamboo lemur P450 genes to be impacted by their hyper-specialized diet.

For this project, we hypothesized that the highly specialized diet observed for bamboo lemurs should be associated with a narrower P450 repertoire, streamlined to those subfamilies that metabolize xenobiotic compounds found in bamboo. Therefore, we predicted that, compared to other generalist Malagasy strepsirrhines, the bamboo lemur species *Prol. simus* and *H. griseus* should exhibit a clear pattern of cytochrome P450 (1) gene loss and (2) signatures of relaxed natural selection, because the specialist diet contains a narrow set of PSMs, of which the primary toxin (*i.e.*, cyanide), is not even metabolized by P450s. Similar to previous work (Chaney et al., 2020), we focus on P450 genes in the *CYP1-3* subfamilies. Briefly, this decision was made because of these subfamilies’ monophyletic placement within the cytochrome P450 superfamily of genes (Kirischian et al., 2011) and the presence of genes encoding both xenobiotic and endobiotic metabolizers within that gene-clade (see Guengerich, 2015).

## 2. Materials and Methods

### 2.1 Generation of *Hapalemur griseus* genome assembly

For this study, we constructed a genome assembly for a male *H. griseus* (named Beavis) housed at the Duke Lemur Center in Durham, North Carolina. A fresh whole blood sample was shipped to the Molecular Primatology Laboratory of New York University for processing and sequencing. High molecular weight (HMW) DNA was extracted from 200 μL of whole blood using a MagAttract HMW DNA Kit (QIAGEN Sciences Inc., Germantown, MD) following the manufacturer’s protocol. DNA was eluted into a low-bind 1.5 mL tube and stored at 4°C prior to size selection. In order to obtain DNA fragments roughly 40 kilobases or longer, the extracted HMW DNA was run through a Blue Pippin DNA Size Selection System (Sage Science Inc., Beverly, MA) using a 0.75% agarose gel cassette with the R1 size standard. Following this size selection protocol, the HMW DNA concentration was quantified using a Qubit 2.0 fluorometer (Thermo Fisher Scientific Inc., Carlsbad, CA) using the broad range standard. The extraction yielded 3.2 ng/μL of HMW DNA.

After extraction and quantification, the HMW DNA sample was sent to NYU Langone Medical School’s Genome Technology Center for quality control tests, which consisted of DNA quantification and fragment size determination using a Tape Station (Agilent Technologies Inc., Santa Clara, CA). After passing this checkpoint, the HMW DNA was prepared for sequencing using a Chromium Genome Linked Reads Kit (10X Genomics Inc., Pleasanton, CA) and sequenced on a HiSeq 2500 DNA Sequencing Platform (Illumina Inc., San Diego, CA) with 1200M reads. The linked-read output data were assembled using the Supernova assembler (Weisenfeld et al., 2017). We used the pseudohaplotype output as our genome assembly for all downstream steps (Table 1).

**Table 1:**
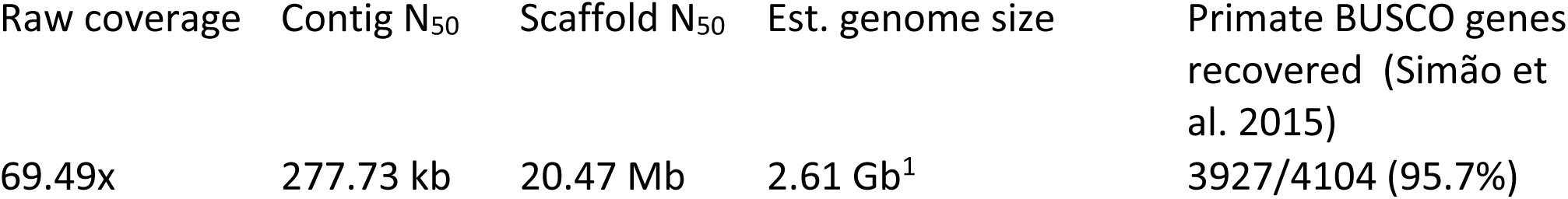
Select statistics from *de novo* assembly of *Hapalemur griseus* (Beavis) using the Supernova assembler.

2.2 Additional data gathering

For both the gene-birth/death and selection-analysis portions of this study, we gathered data from publicly available genome assemblies for 14 species: *Prolemur simus* (Hawkins et al., 2018), *Lemur catta* (Palmada-Flores et al., 2022), *Eulemur flavifrons* and *E. macaco* (Meyer et al. 2015), *Propithecus coquereli* (Lowe and Eddy 1997; Guevara et al. 2021), *Indri indri* (accession number: GCA_004363605.1), *Daubentonia madagascariensis* (accession number: GCA_004027145.1), *Mirza coquereli* (accession number: GCA_004024645.1), *Mirza zaza* (Hunnicutt et al., 2020), and *Microcebus murinus* (Averdam et al., 2011; Lecompte et al., 2016), as well as the following additional species of mouse lemur: *Mic. griseorufus, Mic. mittermeieri, Mic. ravelobensis,* and *Mic. tavaratra* (Hunnicutt et al., 2020). These assemblies, along with any associated annotation files, were downloaded locally and formatted into BLAST databases within Geneious Prime, version 2022.1.1.

We located the loci for all annotated *CYP1-3* homologs in the *L. catta, Prop. Coquereli,* and *Mic. murinus* by using the associated annotation (GFF3) files for each. We defined these loci by the non-P450 genes that bounded them; therefore, those surrounding genes were used initially as queries for local BLAST searches. In this way, each locus was linked to two searches per species. The three reference genomes listed above were used because they are all members of separate strepsirrhine families (Lemuridae, Indriidae, and Cheirogaleidae, respectively), and they were each therefore used as a starting point to extract the desired *CYP1-3* genes or loci for confamilial species. Ideally, a pair of BLAST searches would return results that included the same scaffold. By locating both BLAST hits on each of these scaffolds, we were able to extract genomic regions that were hypothetically orthologous to those P450 loci in the *L. catta* assembly. After locating the scaffolds in each assembly corresponding to each P450 locus, we used LASTZ (Harris, 2007) to interrogate the homology of those scaffolds by aligning them to the confirmed P450 locus from the appropriate confamilial reference genome. Positive results from these alignments were checked using the Mauve genome aligner (Darling et al., 2004) on the same sequences. If output from both of these aligners indicated that the reference had homology with the query scaffold(s), then the annotations from the reference genome were used to extract the corresponding sequence in the other species’ genome. In this way, we mined the genome assemblies listed above for as many complete P450 genes loci as we could confidently locate.

To augment our dataset of codon sequences specifically for our selection analyses (see below), we also devised a *blastn-*based strategy to gather such data from 25 recently published assemblies in the Primate Genome Diversity Project (PGDP; Kuderna et al., 2023). Similar to locating complete P450 loci for the birth-death analysis, we used one reference species for each of three strepsirrhine families comprised by the portion of the PGDP data that we utilized: Cheirogaleidae (reference species: *Mic. murinus*), Indriidae (*Prop. Coquereli*), and Lemuridae *(L. catta*). Each of the *CYP1-3* genes for these references was transposed into a multifasta file with each codon separated by sequence headers. Each of these exons were then used to independently query each of the selected PGDP genome assemblies, which resulted in a large dataset of BLAST hits for each of these searches. For each set of results, any orthologous sequence was used only if it met the following criteria: all of the resultant exons were found on the same contig, these exons were in the same relative order as in the reference (after accounting for strandedness), and this locus did not appear elsewhere in the dataset (after all sequence data had eventually been gathered). In order to maximize our statistical power in the downstream selection analysis, we accepted partial sequences, or sequences with incomplete coverage of each query sequence. In order to construct the trees needed for these analyses, we downloaded all sequence between the first and last exons for each set of BLAST results and used this for alignment and phylogenetics (see below).

### 2.3 Inference of gene birth and death

For this first portion of the study, we used only species for which each *CYP1-3* locus could be wholly collected from a single scaffold or reasonably reconstructed if not found on a single scaffold using the process described above. In order to model the events of gene birth and death in this subset of lemur species, our alignment strategy followed a similar workflow as outlined in previous work with other datasets (Chaney et al., 2018, 2020), but several modifications were made for this project in order to allow for more standardization and automation across subfamilies. First, all of the P450 genes were extracted from each species’ locus according to the annotation file associated with its confamilial reference. Then, all of the genes from a given P450 subfamily were aligned using MAFFT (Katoh & Standley, 2013), and the resulting alignment was stripped of all sites containing any gaps using trimAl (Capella-Gutierrez et al., 2009). After the best-fitting nucleotide substitution model was inferred by jModelTest (Darriba et al., 2012), this stripped alignment was visualized with PhyML 3.0 and the strength of that resulting phylogenetic tree was tested by comparing it to 1000 bootstrap replicates (Guindon et al., 2010).

The gene trees constructed with PhyML were then passed to Possvm (Grau-Bové & Sebé-Pedrós, 2021). This program uses the intrinsic information contained in a phylogram to infer speciation and gene-duplication events; it does this using the species-overlap algorithm in the ETE3 toolkit (Huerta-Cepas et al., 2007, 2016). Briefly, this algorithm compares the intersection of species present in both descendants of an internal node of a tree to the union of species present in those descendants; using these values, the algorithm computes a species-overlap score which it then uses to identify each internal node as either a speciation event, having an overlap score, or a duplication event, having a high overlap score (Huerta-Cepas et al., 2007). Once the identities of each node were estimated in this way, we then manually examined each subtree rooted by a node called as a duplication event to infer whether any gene loss had occurred. This was examined on a case-by-case basis using the reasoning that, after a duplication event, each descendant of that node should recapitulate the organismal phylogeny present at the time of duplication. Therefore, any species missing in one of those subtrees must have lost one of the duplicates born in the earlier duplication event as long as the subtree in question was well-resolved in terms of bootstrap support. In cases where multiple species lineages may be absent, we deferred to the parsimonious hypothesis that a loss event would have occurred prior to the divergence of those lineages, rather than a more complicated hypothesis that the same paralog had been independently lost in both species after their split. We visualized the Possvm output using the program Treerecs (Comte et al., 2020) and then, in some cases, manually modified the depicted gene-evolution scenario in order to accommodate the Possvm results.

### 2.4 Selection analysis

We used primarily used HyPhy’s RELAX program (Wertheim et al., 2015) to test our hypothesis concerning the relaxing of selectional pressure on the P450s covered in this project, but the programs aBSREL (Smith et al., 2018) and BUSTED (Murrell et al., 2015) were also used only in the case of *CYP2C* because of some preliminary tests indicating possible diversifying selection in this gene subfamily (see below: section 3.2). For all runs, we made sure to include estimates of synonymous rate variation because ignoring this parameter and setting it to 1.00, as is often done for many *dN/dS* analyses, has been shown to bias results and, in the case of detecting diversifying selection, inflate the number of false positives (Wisotsky et al., 2021).Each HyPhy run requires, first, a codon alignment stripped of any stop codons and, second, a phylogenetic tree that contains labels for the branches on which *a priori* relaxation of natural selection is hypothesized to have occurred. For the codon alignments, we used preexisting annotations from each of the annotated genome assemblies to extract the likely coding DNA sequence (CDS) from each of the ostensibly complete P450 genes gathered in the first portion of the study. Those annotations were then leveraged to find the likely CDS in unannotated homologous sequences from confamilial relatives.

We aligned the CDS from each subfamily by translation using MACSE (Ranwez et al., 2018), and we then checked each alignment by eye using MEGA 11 (Tamura et al., 2021). Once the alignment had been checked by eye and the maximum number of useable sequences were included, it was exported in FASTA format. The trees used for all HyPhy runs were were built from genomic (*i.e.,* combined exonic and intronic) sequence using MAFFT (Katoh & Standley, 2013). We compared the rows in each subfamily’s final CDS alignment to the leaves of the gene tree constructed earlier with an R script using functions from the *phytools* (Revell, 2012) and *seqinr* (Charif & Lobry, 2007) R packages, and a new tree was written that contained only the intersection of these two sets. This filtered gene tree was then viewed with the web-based *phylotree.js* platform (Shank et al., 2018), which allowed us to easily label branches on each tree. Only foreground branches were labeled for *Prol. simus, H. griseus*, and any internal branches exclusive to the clade formed by these two species. Accordingly, all unlabeled branches were used for the estimation of the background *dN*/*dS* ratios for RELAX. Aside from manually reading the text-file output, the HyPhy Vision platform (http://vision.hyphy.org/RELAX) was used to generate some of the graphs made from the output JSON files. For ancestral-sequence reconstruction of *CYP2F*, we used a HyPhy batch file that makes use of parameters estimated from a selection analysis run beforehand (https://github.com/veg/hyphy-analyses/tree/master/AncestralSequences). Rather than using some other software for ancestral-sequence reconstruction, our use of this HyPhy batch file increased our confidence in the comparability of our RELAX runs and the ancestral sequences used in our *CYP2F* post-hoc analysis. Using these ancestral sequences, we then computed pairwise *dN/dS* values using the Nei-Gojobori (1986)Nei-Gojobori (1986) method to more closely examine signals of neutral evolution or selectional relaxation revealed by the earlier HyPhy RELAX analysis.

## 3. Results

The evidence presented here largely displays a trend in which the common lineage of bamboo lemurs has a diminished or expected number of gene copies relative to generalist species, with several xenobiotic-metabolizing P450 subfamilies exhibiting a resolved gene-tree topology indicating genic loss in one or both of the bamboo lemur species included here.

Because of the other species included in this data set, there are also some insights to be gained from examining other portions of the gene trees outside of the portions focused on our species of interest.

### 3.1 Heightened gene loss within the bamboo-lemur lineage

The topologies of the *CYP1A* and *CYP1B* subfamilies show strong bootstrap support and near-perfect correspondence to the species phylogeny for their member orthologs (Figs. S1 & S2). The only exception to this derives from the *Propithecus coquereli* lineage, which saw the duplication of the *CYP1A1* ortholog after the divergence of the *Prop. coquereli* and *Indri indri* lineages.

The shape of the *CYP2A* gene tree implies substantially more genic turnover than either of the *CYP1* subfamilies (Fig. 1, top-left; Fig. S3). While the *Microcebus murinus* lineage shows a very straightforward trajectory of no duplications, all other species have either duplicated or lost orthologs according to the pattern of relatedness displayed in this tree. Due to the mid-point rooting of this tree by Possvm, it is unclear whether two or three orthogroups are truly present in this case (see Fig. S3), but we opt for a parsimonious interpretation of two orthogroups here (see Fig. 1). Nonetheless, at least one duplication was certainly specific to the *Prop. coquereli* lineage, depicted here by the common node of *CYP2A13L1* and *CYP2A13L2*. The lemurid portion of the tree indicates a gene duplication in the common ancestor of all three species and another duplication that was specific to *L. catta*. Conversely, the topology displayed here implies genic loss for *E. macaco* and for *Prol. simus*.

The only members of *CYP2B* examined in this study are *CYP2B6* homologs because another group of orthologs in the *L. catta* genome assembly (*CYP2B11*) did not show a CDS length that was remotely close to the normal length of a functional primate P450 gene (*i.e.*, *ca.* 1500 nt). In the *CYP2B6* tree (Fig. 1, bottom-left; Fig. S4), we find two major orthogroups among the five species analyzed. One of these shows a topology that, in itself, appears to recapitulate the organismal phylogeny with no duplications. However, the other orthogroup exhibits a signature of interesting dynamism. There are two serial duplication events in the *Prop. coquereli* lineage (*CYP2B6L1, 3,* and *4P*). In the lemurid portion of this subtree, there are two more serial duplications prior to the speciation of *L. catta* and *Prol. simus.* The pattern in this part of the tree most parsimoniously indicates, first, that the bamboo-lemur common ancestor lost genes orthologous to the *CYP2B6L3* and *CYP2B6L4* genes in the *L. catta* assembly and, second, that the *H. griseus* lineage further lost its final remaining gene in this orthogroup.

The *CYP2C* genes yielded a phylogeny whose complexity is underscored by the large number of duplications within it (Fig. 1, right; Fig. S5). Possvm separated this phylogeny into five coherent orthogroups. The first of these (*i.e.,* OG-0) contained a duplication prior to the divergence of *L. catta* and the bamboo lemurs. Following the *Prolemur-Hapalemur* divergence, *Prol. simus* lost one of these genes but the other paralog was lost by *H. griseus*; thus, both species retained separate paralogs. OG-1 and OG-2 lack *Prol. simus* genes but do contain genes belonging to *H. griseus*; this pattern indicates two lineage specific events of gene loss for *Prol. simus.* While this latter species, the greater bamboo lemur, showed a pattern of complete loss for these orthogroups, *Mic. murinus* experienced a duplication in OG-2. All lemurid genes were ostensibly lost from OG-3, while the mouse-lemur lineage saw a single-gene expansion of this orthogroup. The final clade of genes, OG-4, contained three duplication events and, parsimoniously, three instances of gene loss. While *L. catta* retained every one of the duplicated genes from this sequence of events, bamboo lemurs likely lost the ortholog to *L. catta CYP2C18L1*, and both *Prol. simus* and *H. griseus* further lost one more gene each.

However, these two final losses were lineage-specific, and each involved a separate genic lineage. Thus, with the exception of genes in one orthogroup (*i.e.*, OG-3), bamboo lemurs exhibited all of the losses detected in this tree.

Similar to *CYP2C*, the *CYP2D* subfamily also showed an increased amount of evolutionary dynamism with regard to gene birth and death (Fig. 2, top-left; Fig. S6). Along a basal branch of the phylogeny, there are two serial duplications, resulting in three *Mic. murinus* homologs equally related to all lemurid *CYP2D* homologs. Each of the black lemur (*E. macaco*) species has two very similar *CYP2D* genes which arose from the duplication specific to the *Eulemur* lineage. Prior to the divergence of the *L. catta* lineage from that of the bamboo lemurs, there is some uncertainty in these results whether only a single duplication occurred or whether two occurred. This uncertainty stems from two nodes displaying low bootstrap support (73.0% and 57.5%; see Fig. S6) in the original PhyML tree. The topology displayed in Figure S6 describes two duplications and parsimoniously implies just as many deletions. In light of the low bootstrap support for this pattern, however, it is conceivable that the true topology could place the *L. catta CYP2D17L2* gene into the clade with the two bamboo-lemur genes. Such a revision would reduce the number of deletions and duplications. The other duplicated gene in the lemurid lineage, while secondarily duplicated in the *L. catta* lineage, was lost in the *H. griseus* lineage.

Lastly, no functional homologs for the *CYP2D* subfamily could be retrieved for *H. griseus* because no open reading frames devoid of stop codons could be found in the extracted sequence for this species’ single *CYP2D* gene. However, this isolated instance may be due to an assembly artifact, as the *CYP2D* locus was located on the ends of two separate scaffolds in the original Supernova assembly. In all, the general pattern of this subfamily is similar to that of *CYP2C*: evidence of heightened genic turnover with an emphasis on loss in the specific lineage(s) of the bamboo lemurs.

The *CYP2J* subfamily divided into three orthogroups (Fig. 2, top-right; Fig. S7). The first of these (OG-0) contains only a single duplication node before the speciation node for the split of *L. catta* from the bamboo lemurs. The complete lack of any bamboo-lemur genes in this portion of the tree parsimoniously indicates a loss of that duplicate prior to the *Prolemur-Hapalemur* divergence. The remaining two *CYP2J* orthogroups (OG-1 and OG-2) arose as the product of a duplication event prior to the first speciation event of all species involved in this analysis. While no more duplications were called in the OG1-OG2 clade, several deletions are implied by the tree’s topology. First, one gene appears to have been lost in the *Prop. coquereli* lineage. Second, another deletion was ostensibly lost in the ancestral lemurid lineage. Last, *H. griseus* lost one more homolog after that lineage’s split from that of *Prol. simus.* Thus, the *CYP2J* subfamily displays more of the above-mentioned pattern wherein the bamboo lemurs stand out for their likelihood of losing P450 genes.

The topologies of the remaining *CYP2* subfamilies (*i.e.*, *E, F, G, R, S, U,* and *W*) show no evidence of duplication (Figs. S8-S13). Degeneration of one *CYP2F* gene may have occurred in the *Mic. murinus* lineage (see Fig. S9), but anthropoids and at least some lorisiforms appear to possess only a single *CYP2F* gene as well (Chaney et al., 2020; Chaney, 2023). Therefore, the ancestral primate number of *CYP2F* orthologs is uncertain without the inclusion of a phylogenetically broader set of species. With this single possible exception, bamboo lemurs have clearly maintained the same number of gene copies in each of these subfamilies as the other lemur species analyzed here.

The final subfamily of this part of the analysis further underscores the pattern described for other subfamilies above (Fig. 2, bottom-left; Fig. S14). While OG-0 within the *CYP3A* tree possesses only speciation nodes, one loss is evident in the absence of *Prop. coquereli* from this clade. The other part of the tree exhibits a very complex topology because of the number of duplication nodes contained within it. Many of these outside of the lemurid portion of the tree are taxon-specific: *Prop. coquereli* and *Mic. murinus* each experienced two and four exclusive duplication events, respectively. The lemurid portion of OG-1 shows at least three duplication events prior to the divergence of *L. catta* and from the ancestral bamboo-lemur lineage. One more duplication event, leading to the *L. catta* genes *CYP3A4L1* and *CYP3A4L2*, would be most parsimoniously interpreted as having occurred specifically in ring-tail lemur lineage, but it is not ruled out that this duplication may have occurred in an ancestral species and been lost by both bamboo lemur species. Regardless, at least two separate events of genic loss seem to have occurred in the lineage of the bamboo lemurs. Thus, while *L. catta* appears to exhibit six separate *CYP3A* homologs, bamboo lemurs possess half that number; furthermore, we conclude that this pattern is attributable nearly wholly to gene loss by bamboo lemurs rather than lineage-specific expansion of *CYP3A* for *L. catta*.

### 3.2 Selection relaxed on two P450 xenobiotic metabolizers

Out of all P450 subfamilies studied here, only *CYP2F* and *CYP2J* showed significant signals of selectional relaxation within the bamboo-lemur clade (Table S2). The *CYP2F* subfamily’s estimated *dN/dS* distributions were interesting in that all rate classes in the test-branch distribution collapsed to a ratio of 1.00, indicating neutral evolution of these genes since their divergence from their *L. catta* orthologs. After examining all bamboo-lemur sequences and comparing them to their ancestral reconstructions, we found that all *dN* – *dS* calculations for the bamboo-lemur *CYP2F1* and *CYP2F2* ancestral sequences and all of their descendent sequences were virtually zero (*CYP2F1* range: -9.2e-3, -9.4e-4; *CYP2F2* range: 0.018, -6.6e-3).

This result is consistent with the above RELAX results, which indicate that both *CYP2F* orthologs appear to be evolving neutrally in the bamboo-lemur clade.

The *CYP2C* subfamily exhibited a significant signal of selectional *intensification* (*p* = 6.8e-8, *k* = 1.42; see Table S3). Examination of the omega distribution for the test branches showed that this appears to be driven by a heightened estimate for the highest *dN/dS* rate class, and follow-up analysis with aBSREL and BUSTED showed this result was driven by a site in the *H. griseus* ortholog for *CYP2C21* (Fig. S15). Indeed, aBSREL detected diversifying selection only along the this terminal branch (likelihood ratio = 41.92, *p* < 0.00001), with a very high estimate for the highest *dN/dS* rate class (> 1000). Some, but not all, of this variation was shared by the closely related *H. meridionalis*, but this species’ terminal branch did not show any positive result from our aBSREL or BUSTED analyses.

## 4. Discussion

In this study, we have demonstrated that the genomes of two bamboo lemur species (*H. griseus* and *Prol. simus*) harbor, at a conservative maximum, the modal number of cytochrome P450 genes across the other lemur species included here (Table S1). Indeed, in most cases where losses of P450 genes were inferred, they are most parsimoniously attributed to a bamboo lemur-specific branch of the phylogeny in question (see Figs. 1-2). Thus, bamboo lemurs appear to show a narrower repertoire of xenobiotic-metabolizing P450s insofar as they have generally fewer genes than other species of lemur, especially in comparison to their closest relative, *Lemur catta.* Bamboo lemurs also show signals of relaxation of natural selection for the *CYP2F* and *CYP2J* subfamilies (see Fig. 3), which are known to be involved in xenobiotic metabolism (Danielson, 2002; Guengerich, 2015).

These results are perhaps best understood in terms of an evolutionary tradeoff on detoxification strategies (Dearing et al., 2005; Freeland & Janzen, 1974; Ueno, 2001). In a pivotal paper outlining several nutritional-ecological hypotheses, Freeland and Janzen (1974) formalized the notion that generalist and specialist herbivores should differ inversely with regard to the breadth of their arsenals for the metabolism of toxic plant secondary metabolites. In their view, herbivores should possess a comparatively wide array of detoxification mechanisms to handle low doses of a commensurately broad variety of toxins, and specialist herbivores should, conversely, possess a smaller array of such mechanisms able to handle large doses of a narrow variety of toxins (Dearing et al., 2005; Freeland & Janzen, 1974; Marsh et al., 2006). The general pattern of copy-number differences and the more limited result of selectional relaxation shown in this study seems to support this hypothesis in the case of bamboo lemurs, alongside this taxon’s high degree of specialization (see Table S1) and the stupendous amounts of cyanide consumed by them (Ballhorn et al., 2009; Glander et al., 1989).

In this set of results, the *CYP2F* subfamily stood out from other P450 subfamilies by having a particular signal of selectional relaxation that suggested neutral evolution, as all of the rate classes for *CYP2F* bamboo-lemur orthologs collapsed toward *dN/dS* = 1.00. Follow-up comparisons of *dN* and *dS* were consistent with this result (Table S4). Although this P450 subfamily is most highly expressed in the respiratory tract, hepatic expression has been demonstrated, albeit at lower levels (Duff et al., 2015; Fagerberg et al., 2014; Yue et al., 2014). Human CYP2F1 and its model-rodent homologs can activate coumarins through epoxidation reactions, leading to cytotoxicity and necrosis at varying species-dependent levels (Born et al, 2002; Vassalo et al., 2004). Similarly, CYP2F enzymes have been linked with bioactivation of the toxin naphthalene, also through an oxidation reaction (Lanza et al., 1999; Li et al., 2011; Zhang & Ding, 2008). These biochemical associations are notable here because chromatographic assays and microbiome studies of various species of bamboo have implied or directly indicated the presence of both naphthalene (Li et al., 2002; Takahashi et al., 2010) and coumarins (Coffie et al., 2014) in those bamboo species (reviewed by Gagliano et al., 2022). Generalizing from these findings, we would hypothesize, first, that naphthalene and coumarins are present in Malagasy bamboo consumed by bamboo lemurs and, second, that bamboo lemurs would be susceptible to toxicity from their metabolite oxides (napthalene: see Yost et al., 2021; coumarins: see Foroozesh et al., 2019). Therefore, neutral evolution of bamboo lemurs’ *CYP2F* orthologs may have allowed for marginally lowered efficiency of the bioactivation of these PSMs into more toxic metabolites. Experimental testing of this hypothesis would help to clarify this interesting result.

*CYP2J,* the other subfamily to show significant relaxation of selection pressure for bamboo lemurs, is largely responsible for the metabolism of arachidonic acid, an endogenous molecule, and other polyunsaturated fatty acids (Xu et al., 2013; Q.-Y. Zhang & Ding, 2008).

However, CYP2J enzymes have been shown to be important in the metabolism of some xenobiotic compounds such as linoleic acid and a variety of pharmaceutical compounds (Capdevila & Wang, 2013; Falck et al., 2009; Hildreth et al., 2020; Michaelis et al., 2003; Pidkovka et al., 2013; Tang et al., 2009; M.-Z. Zhang et al., 2013; cited in Guengerich, 2015). We suggest therefore that bamboo-lemur diets are likely sparse in such xenobiotics that would be targeted by CYP2J enzymes.

The difference between bamboo lemurs and the sifaka *Propithecus coquereli* stands out as a second confirmatory signal of Freeland and Janzen’s (1974) tradeoff hypothesis because, while both of these taxa are generally classified as folivorous, they differ in their amount of dietary specialization. As mentioned previously, all species of bamboo lemur that have been studied are heavily reliant on either grasses (usually bamboo) or sedges, and very few plants, often only a single species, are required to describe a large majority of their diets (see Table S1). Conversely, sifakas (*Propithecus* spp.) consume much more varied foods (Hemingway, 1998; Norscia et al., 2006; Powzyk & Mowry, 2003; Rowe et al., 2016), seasonally trending toward gramnivory or frugivory depending on the species in question (Irwin 2006). Despite this dietary diversity, sifakas do possess clear gastrointestinal adaptations related to hindgut fermentation, such as a greatly elongated intestine and an expanded, haustrated cecum (Campbell et al., 2000). For this reason, sifakas are often described as *anatomical folivores*, with the implication that they are more behaviorally flexible with their diet choice than one might be led to believe upon viewing their gastrointestinal anatomy. Meanwhile, bamboo lemurs of all three major species possess a relatively simpler gastrointestinal anatomy, with an overall relatively shorter intestine and a shorter, though completely haustrated, colon (Campbell et al. 2000; Yamashita et al. 2008). All of this information serves to position sifakas and bamboo lemurs on opposite sides of a specialist-generalist continuum among herbivores.

Such dietary differences between these two taxa helps to clarify an ultimate explanation for the opposite pattern of gene birth and death seen in the *CYP2B6* gene cluster for bamboo lemurs and *Prop. coquereli* (see Fig. 1). This is also true, but to a less pronounced degree, for the *CYP2A* subfamily (see Fig. 1). In both of these cases, bamboo lemurs exhibit clade-specific gene losses and *Prop. coquereli* shows an opposite signal of lineage-specific gene duplications. Indeed, (Guevara et al. 2021), utilizing this same genome assembly for *Prop. coquereli,* independently found that these two P450 subfamilies had been enriched in so-called “rapidly evolving genes,” or genes with increased *dN/dS* ratios according to PAML branch-site models (Yang, 2007). Because both *CYP2A* and *CYP2B* subfamilies are involved in the metabolism of xenobiotic compounds (Guengerich 2015; Orr et al. 2020), at least these two P450 subfamilies stand out as candidate gene groups supporting Freeland and Janzen’s (1974) prediction, that specialist and generalist herbivores should differ oppositely in the diversity of their means for detoxifying plant secondary metabolites. This interesting natural experiment, of comparing bamboo lemurs and sifakas (or indriids in general), deserves future investigation.

A second line of inquiry is the contrast between *Lemur catta* and the bamboo lemurs. The number of xenobiotic-metabolizing P450 genes in the *L. catta* genome was always greater than or equal to those in either bamboo lemur genome. The highly varied and very seasonal diet documented for *L. catta* (Jolly, 1966; LaFleur, 2012; Sauther, 1992; Sauther et al., 1999; Simmen, Sauther, et al., 2006; Soma, 2006) stands out as a potential explanation for this large number of detoxification mechanisms. Indeed, Sauther et al. (1999) argue that the best dietary classification for *L. catta* should be “opportunistic omnivores” (p. 122) because of the breadth of this species’ dietary repertoire and the drastic spikes of certain foods based on their seasonal availability. This species also appears to be relatively inured to plant secondary metabolites based on the high tannin and alkaloid content of their foods (Gould et al., 2009; Simmen, Sauther, et al., 2006) and their relatively high rejection threshold for the bitter alkaloid quinine (Simmen, Peronny, et al., 2006). This constellation of information portrays a pattern of an animal that is exposed to a broad array of plant secondary metabolites and, perhaps because of the annual variety of their foods, exposed to them at relatively low levels.

While *L. catta* possesses a sizeable number of xenobiotic-metabolizing P450 genes in comparison to both species of bamboo lemur (see Table S1), these genes are not the products of species-specific duplications, as is plain from the phylogenetic pattern of relatedness among all the genes falling into the clade of ringtail and bamboo lemurs. Instead, many of these duplicated genes seem to have arisen during the internode of a lemurid common ancestor, or that of a common ancestor of the *Hapalemur* and *Lemur* genera, and then subsequently lost by one or both of the bamboo lemurs species included here. This result accords well with experimental work on bitter taste receptors (TAS2R16), which shows that the species of bamboo lemur included here (as well as *H. aureus*) show a decreased sensitivity to several bitter β-glycoside compounds (Itoigawa et al., 2021). Revealingly, however, the missense mutations that most altered the response of the TAS2R16 protein were separate for *Prol. simus* and *Hapalemur* spp., and ancestral reconstructions of the last common ancestors for both the genus *Hapalemur* and for all bamboo lemurs showed a more sensitive response to these bitter compounds than any of the present-day species-specific TAS2R16 isoforms (Itoigawa et al., 2021). Given this earlier work and the pattern of P450 gene birth and death presented here, a generalist dietary pattern is clearly the best estimate for an ancestral ecology of the *Hapalemur-Lemur* last common ancestor. Furthermore, the complex pattern of loss and retention of separate P450 paralogs by *Prol. simus* and *H. griseus* seems to agree with the conclusion of Itoigawa et al. (2021) that these species’ varying, but in all cases intense, specialization on cyanogenic bamboo may have evolved in parallel rather than what would be predicted based on parsimony, namely that this specialization would have evolved in the common ancestor of all extant bamboo-lemur species.

On a methodological note, the pattern of gene gain and loss reported here is certainly not the conclusion that would be reached by using an approach where the gene-count table is parsimoniously compared to an organismal phylogeny to estimate ancestral states of gene counts, as with the commonly used program for studying gene-family evolution, CAFE (De Bie et al., 2006). This software program takes phylogeny into account only insofar as the species themselves are concerned but does not include any information about the genes’ own phylogenetic relatedness, and for this reason, we see it as an undesirable alternative to examining a phylogeny built from the genes themselves. While CAFE is very useful in its ability to be easily automated, manual examinations of gene trees, or perhaps the use of programs such as Possvm (Grau-Bové & Sebé-Pedrós, 2021) that make direct use of gene trees via the species-overlap method (Huerta-Cepas, et al., 2007, 2016), would yield more accurate ancestral reconstructions of gene copy-number differences among species.

A lingering question remains as to what compounds, or families of compounds, are leading to the evolutionary expansions and contractions on display in this study. However, there is currently a clear disconnect in the literature that inhibits any discussion between those who have precise knowledge of P450 substrate compounds (*e.g.,* Guengerich 2015) and those who perform phytochemical assays on foods of wild primates (*e.g.,* Eppley et al. 2017; Thurau et al. 2021; Windley et al. 2022). The disconnect arises from differing levels of scale on which each of these two groups operates: pharmacologists often use precise molecules (*e.g.*, benzo[a]pyrene, aflatoxin B1) in their experiments and parlance, while phytochemists and primatologists speak in the more general terms of compound families (*e.g.*, alkaloids, condensed tannins). This nomenclatural gulf between these two camps is a foggy one, and it serves to cloud any inferences that may be gained about the functions of free-ranging, non-model primate species.

## 5. Conclusion

Here, we have found that bamboo lemurs have, on average, fewer *CYP1-3* gene copies than other lemurs, and this is especially true when they are viewed alongside their closest relative, the ringtail lemur. Viewed another way, bamboo lemurs have, at most, the same number of P450 genes per subfamily as *L. catta*, and they often have fewer such genes. The retention of these specific P450 homologs was not parsimonious when compared with the organismal phylogeny, and the complexity of this pattern may imply some information about bamboo-lemur evolution in general, namely that their specialization on cyanogenic bamboo may have evolved in parallel rather than as a sort of ecological synapomorphy early after the divergence of these two lineages. Only two P450 subfamilies (*CYP2F* and *CYP2J*), both of which are linked with xenobiotic metabolism in humans (Guengerich, 2015), showed a statistical decrease in the intensity of selection on the CDS for the genes within those subfamilies. As in previous work on the evolution of the cytochrome P450 superfamily of genes, this topic provides an interesting and complex window into biological evolution. Future work that includes other species of bamboo lemur (*e.g.*, *H. alaotrensis, H. aureus*) will doubtless only clarify that view.

## Acknowledgments

This project was supported by NSF BCS1919857 to AJT and MEC and NSF BCS1717188/BCS1718715/BCS1718339 to CMB and AJT. Research reported in this publication was supported by the National Center For Advancing Translational Sciences of the National Institutes of Health under Award Number TL1TR003019. The content is solely the responsibility of the authors and does not necessarily represent the official views of the National Institutes of Health. Transfer of the *Hapalemur griseus* sample from Duke University to Kent State University was coordinated by Erin Ehmke; this is Duke Lemur Center publication number XXXX. We are thankful to Roy Heath and Joshua Talbott for access to and support with high-power computing at Kent State University. Andrew Burrell and Cody Ruiz were instrumental in extracting the HMW-DNA and preparing the library for sequencing the *H. griseus* genome. MEC is grateful to Caylee Heiremans and Peter Chaney for their support during and beyond this work.

## 6. Author contributions

MEC had the following roles for this study: conceptualization, data curation, formal analysis, investigation, methodology, visualization, and writing (original draft). AJT had the roles of funding acquisition, resources, software, supervision, and writing (reviewing and editing). CMB was responsible for methodology, project administration, resources, supervision, and writing (reviewing and editing).

### Data Accessibility and Benefit-sharing

FASTQ reads from the genomic sequencing of *Hapalemur griseus* have been deposited in the NCBI SRA (accession number XXXXX), and the alignments and trees necessary to reproduce this work are available on Dryad.

---

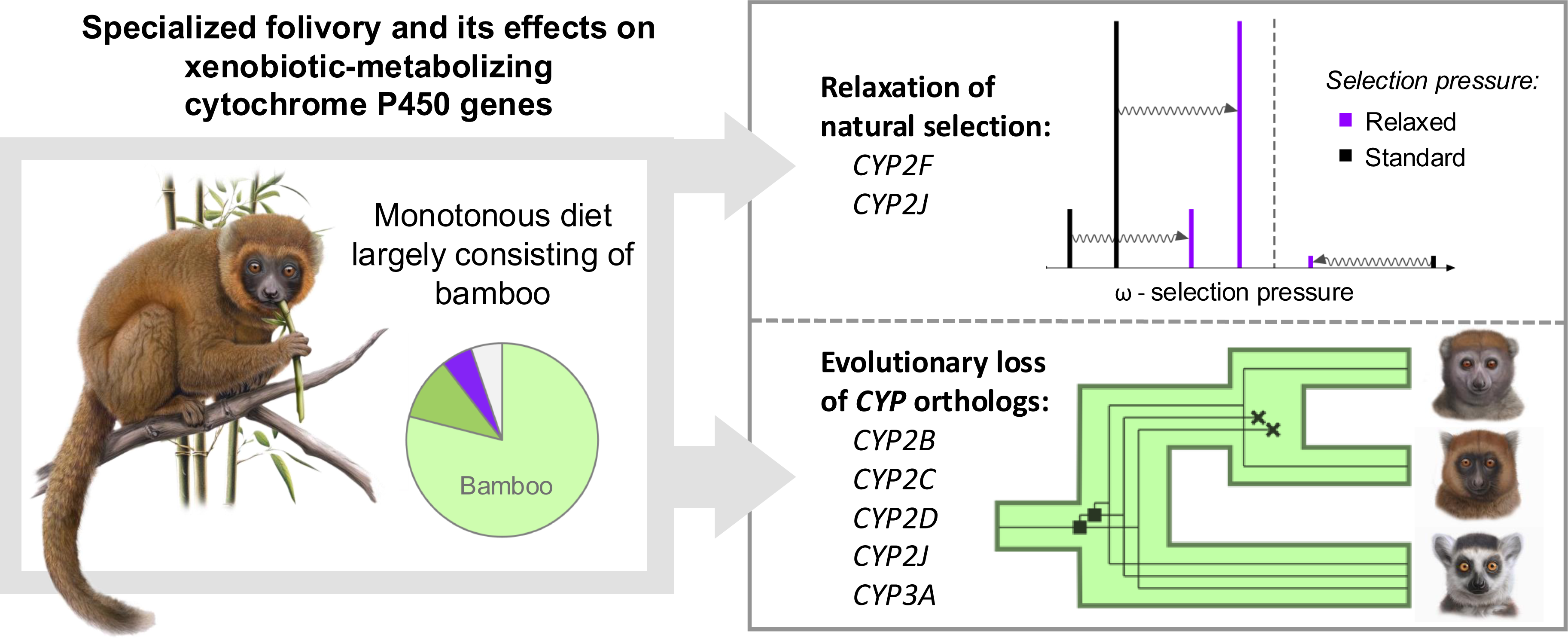

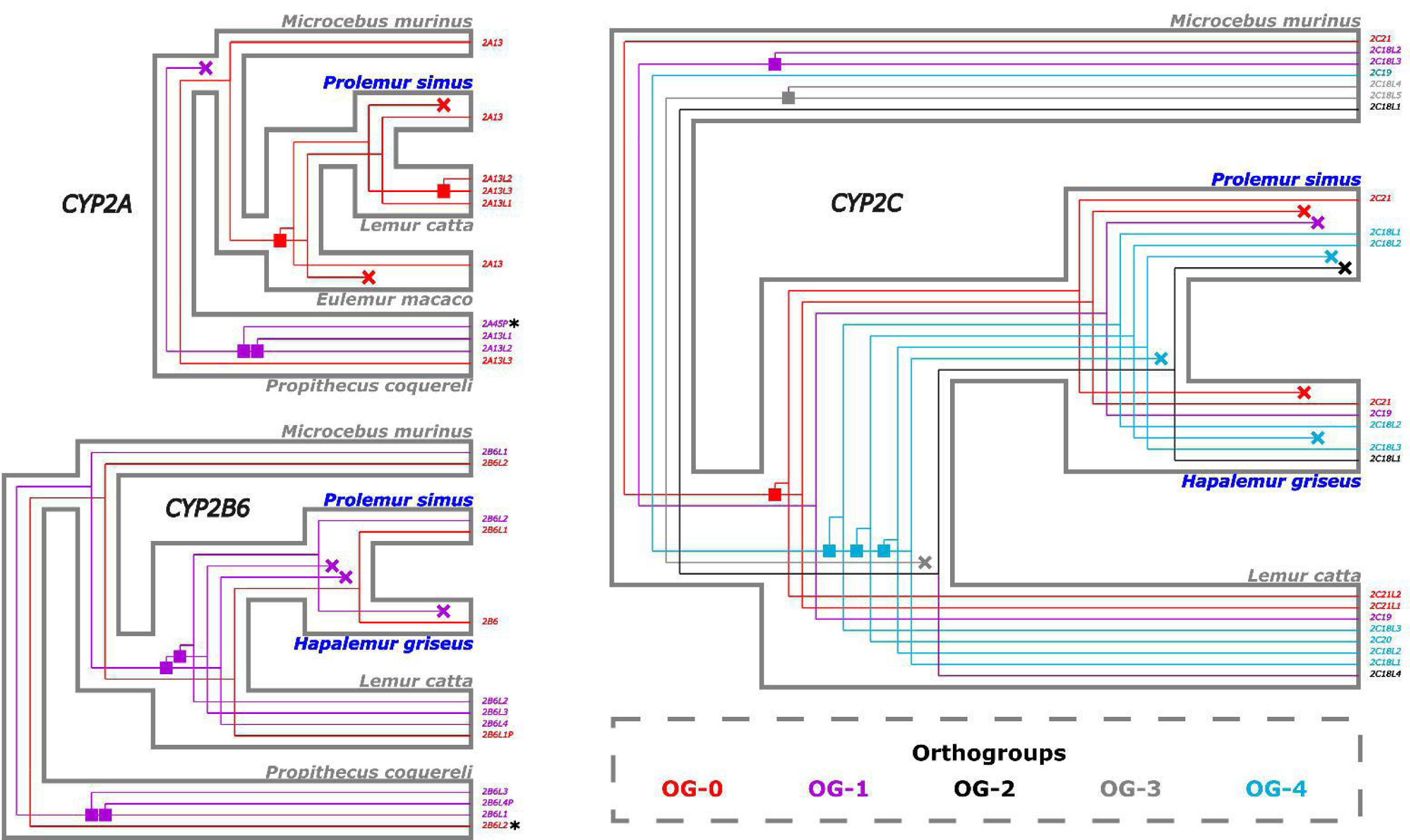

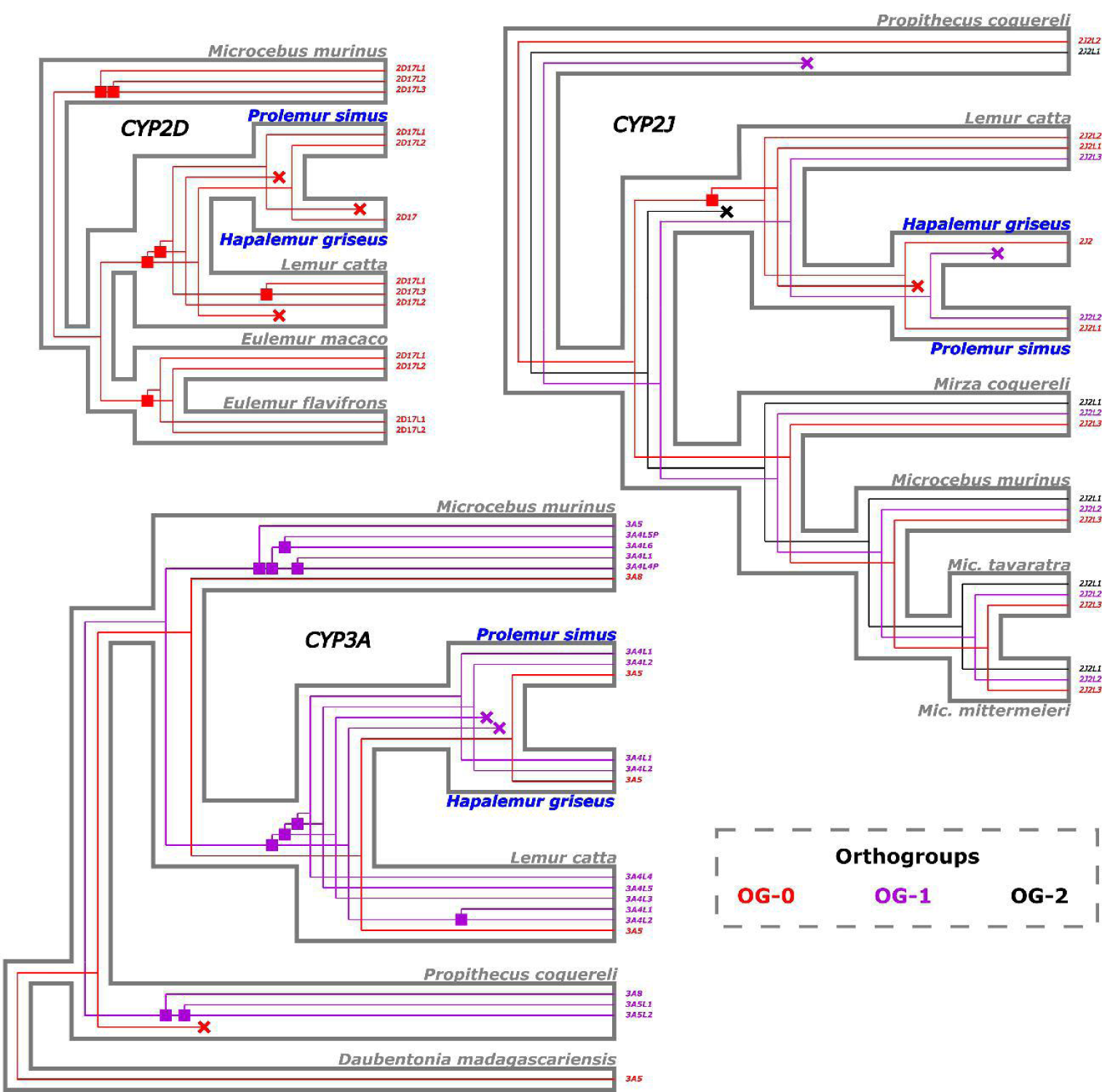

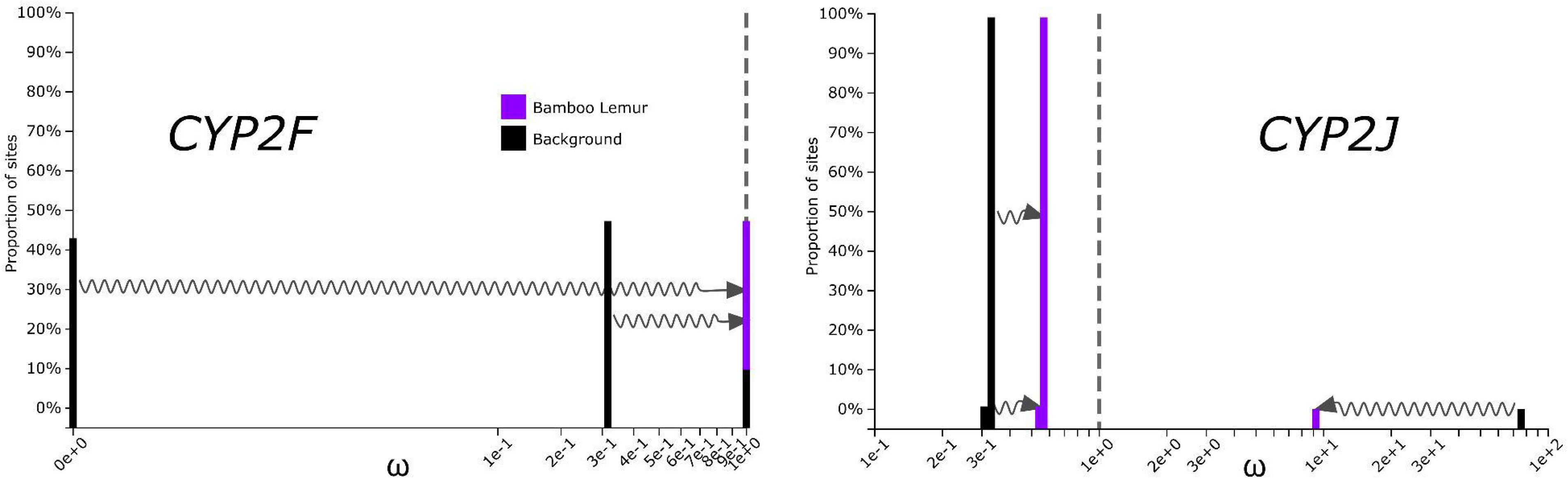

