## Supplementary material for "Hyper-specialized primates possess a reduced suite of xenobiotic-metabolizing cytochrome P450 genes": Online supplement

Table S1: Diets of five bamboo lemur species.

| Species | Study site | Duration of study | Bamboo and grass | Non-grassy foliage | Fruit | Other | References |
| --- | --- | --- | --- | --- | --- | --- | --- |
| *H. alaotrensis* | Lake Alaotra | 15 months | 95.4 | 2.0 |  | 0.3 | Mutschler, 1999 |
| *H. griseus* | Ranomafana National Park (RNP) | 12-24 months | 88 | 5.8 | 5 |  | Tan, 1999 |
| *H. griseus* | RNP | 12 months | 90.7 | 4 | 1.2 | 2.3 | Overdorff et al., 1997 |
| *H. meridionalis* | Mandena Conservation Zone | 62 hours | 76.0 | 21.3 | 1.9 | 0.8 | Eppley et al., 2011 |
| *H. aureus* | RNP | 2 years | 88 | 3 | 4 | 5 | Tan, 1999 |
| *P. simus* | RNP | 2 years | 98 |  | 0.5 | 1.5 | Tan, 1999 |
| *P. simus* | Ambalafary, Brickaville | 112 days | 92.0 | 1.5 | 6.5 |  | Mihaminekena  et al., 2024 |
| *P. simus* | Sahavola, Brickaville | 133 days | 71.7 | 22.7 | 5.3 | 0.3 | Mihaminekena  et al., 2024 |
| *P. simus* | Vatovavy Forest | 193 days | 41.5 | 48.6 | 6.1 | 5.9 | Mihaminekena  et al., 2024 |

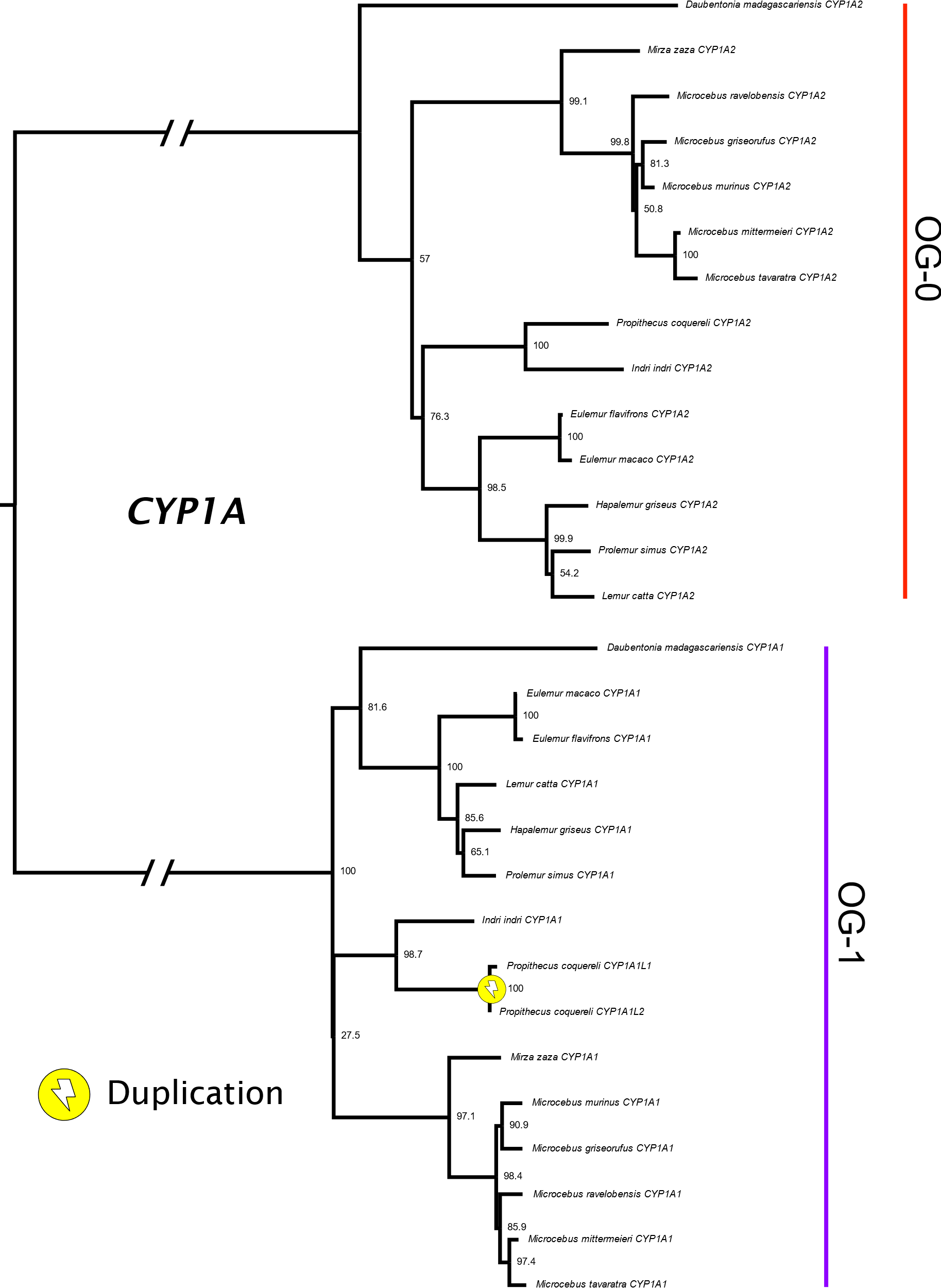

Figure S1: Phylogenetic tree for the *CYP1A* subfamily among 14 species of lemur.

Bootstrap support values are displayed as percentages for each node. In this tree, the branch separating the two orthogroups was particularly long, but it has been abbreviated for the sake of clarity in this figure. The single duplication in this tree was confirmed by Possvm.

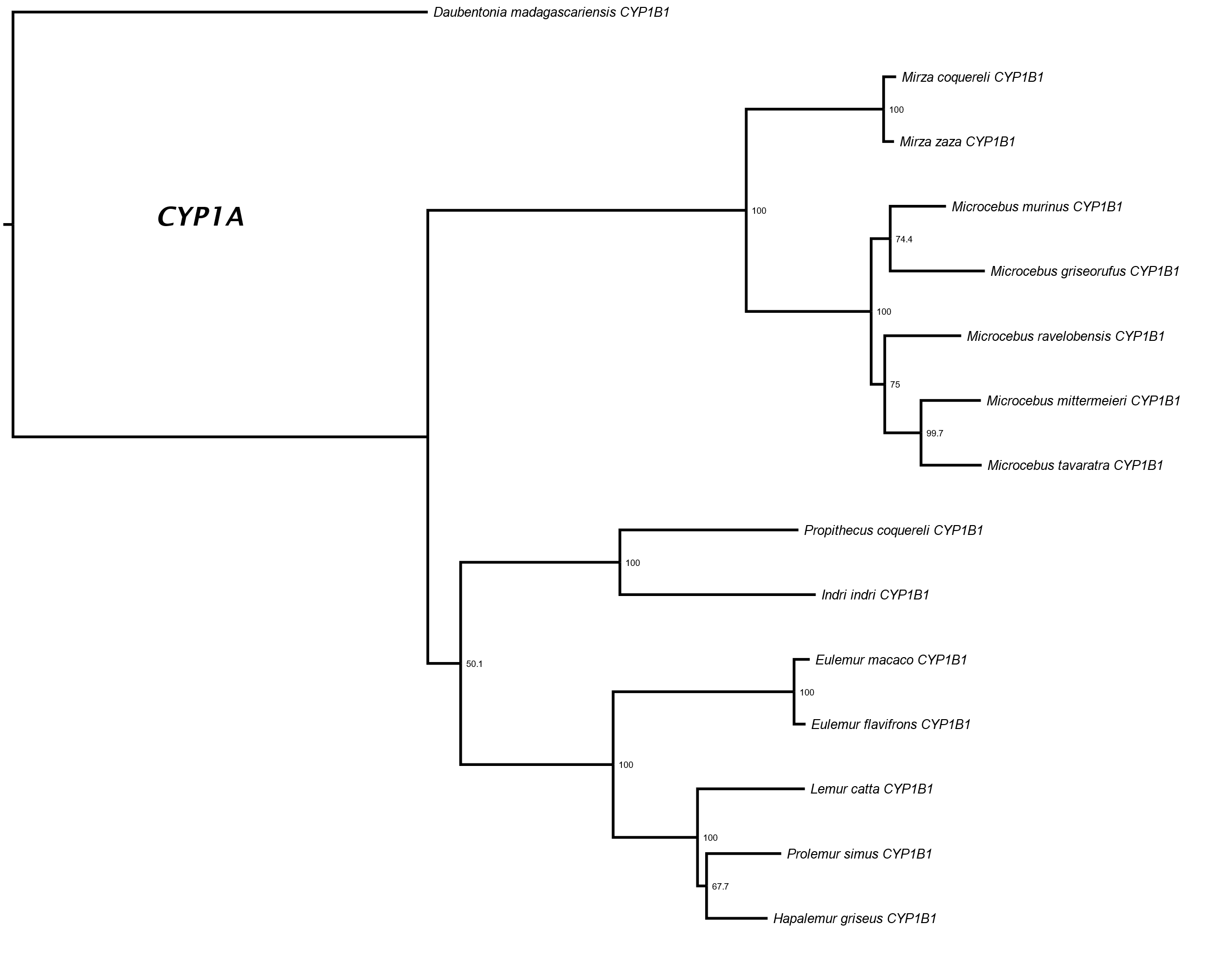

Figure S2: Phylogenetic tree for the *CYP1B* subfamily among 15 species of lemur.

Table S2: Genomic locations of *CYP1-3* loci in the *Lemur catta* genome assembly (mLemCat1; Palmada-Flores et al. 2022).

| P450  Subfamily | Number of Homologs Recovered* | | | | | | | | | | | | | | Bounding Genes in *Lemur catta* Assembly (Searchable NCBI Gene ID) | |
| --- | --- | --- | --- | --- | --- | --- | --- | --- | --- | --- | --- | --- | --- | --- | --- | --- |
|  | **Dm** | **Lc** | **Hg** | **Ps** | **Ef** | **Em** | **Pc** | **Ii** | **Mm** | **Mg** | **Mr** | **Mt** | **Mc** | **Mz** |  |  |
| 1A | 2 | 2 | 2 | 2 | 2 | 2 | 3 | 2 | 2 | 2 | 2 | 2 |  | 2 | *EDC3*  (123624360) | *CSK*  (123624393) |
| 1B | 1 | 1 | 1 | 1 | 1 | 1 | 1 | 1 | 1 | 1 | 1 | 1 | 1 | 1 | *rmdn2*  (123636810) | *ATL2*  (123636813) |
| 2A |  | 3 |  | 1 |  | 1 | 3 |  | 1 |  |  |  |  |  | *AXL*  (123624081) | *EGLN2*  (123624299) |
| 2B6 |  | 3(1) | 1 | 2 |  |  | 3(1) |  | 2 |  |  |  |  |  |  |  |
| 2F |  | 2 | 2 | 2 |  | 2 | 2 |  | 2 |  |  |  |  |  |  |  |
| 2G |  | 1 | 1 | 1 | 1 | 1 | 1 |  | 1 | 1 | 1 | 1 |  |  |  |  |
| 2S |  | 1 | 1 | 1 | 1 | 1 | 1 | 1 | 1 | 1 | 1 |  |  | 1 |  |  |
| 2C |  | 8 | 4 | 3 |  |  |  |  | 7 |  |  |  |  |  | *PDLIM1*  (123650041) | *HELLS*  (123650256) |
| 2D |  | 3 | 1 | 2 | 2 | 2 |  |  |  |  |  |  |  |  | *TCF20*  (123640280) | *NDUFA6*  (123640284) |
| 2E | 1 | 1 | 1 | 1 | 1 | 1 | 1 | 1 | 1 | 1 | 1 | 1 |  | 1 | *SYCE1* (123649855) | *SCART1*  (123650336) |
| 2J |  | 3 | 1 | 2 |  |  | 2 |  | 3 |  |  | 3 | 3 |  | *C3H1orf87*  (123634711) | *HOOK1*  (123635589) |
| 2R |  | 1 | 1 | 1 |  |  | 1 | 1 | 1 | 1 | 1 | 1 |  | 1 | *PDE3B*  (123642476) | *cgrp2l*  (123641653) |
| 2U |  | 1 | 1 | 1 | 1 | 1 | 1 | 1 | 1 | 1 | 1 | 1 | 1 | 1 | *SGMS2*  (123627733) | *HADH*  (123627735) |
| 2W |  | 1 | 1 | 1 | 1 | 1 | 1 | 1 | 1 | 1 | 1 | 1 | 1 | 1 | *cox19*  (123632544) | *C2H7orf50*  (123632548) |
| 3A | 1 | 5 | 3 | 3 |  |  | 2 |  | 4(2) |  |  |  |  |  | *TMEM225B*  (123631586) | *TRIM4*  (123631595) |

For each bounding gene, the NCBI unique identifier is given in parentheses. In cases where the symbols for these genes are not capitalized in this table, the official symbol was a generic one that did not suggest an information about the gene’s homolog (e.g., LOC123638525). These lower-case symbols therefore do not correspond to those in the public annotation file.
*Dm = *Daubentonia madagascariensis*; Lc = *Lemur catta*; Hg = *Hapalemur griseus*; Ps = *Prolemur simus*; Ef = *Eulemur flavifrons*; Em = *E. macaco*; Pc = *Propithecus coquereli*; Ii = *Indri indri*; Mm = *Microcebus murinus*; Mg = *Mic. griseorufus*; Mr = *Mic. ravelobensis*; Mt = *Mic. tavaratra*; Mc = *Mirza coquereli*; Mz = *Mirza zaza*

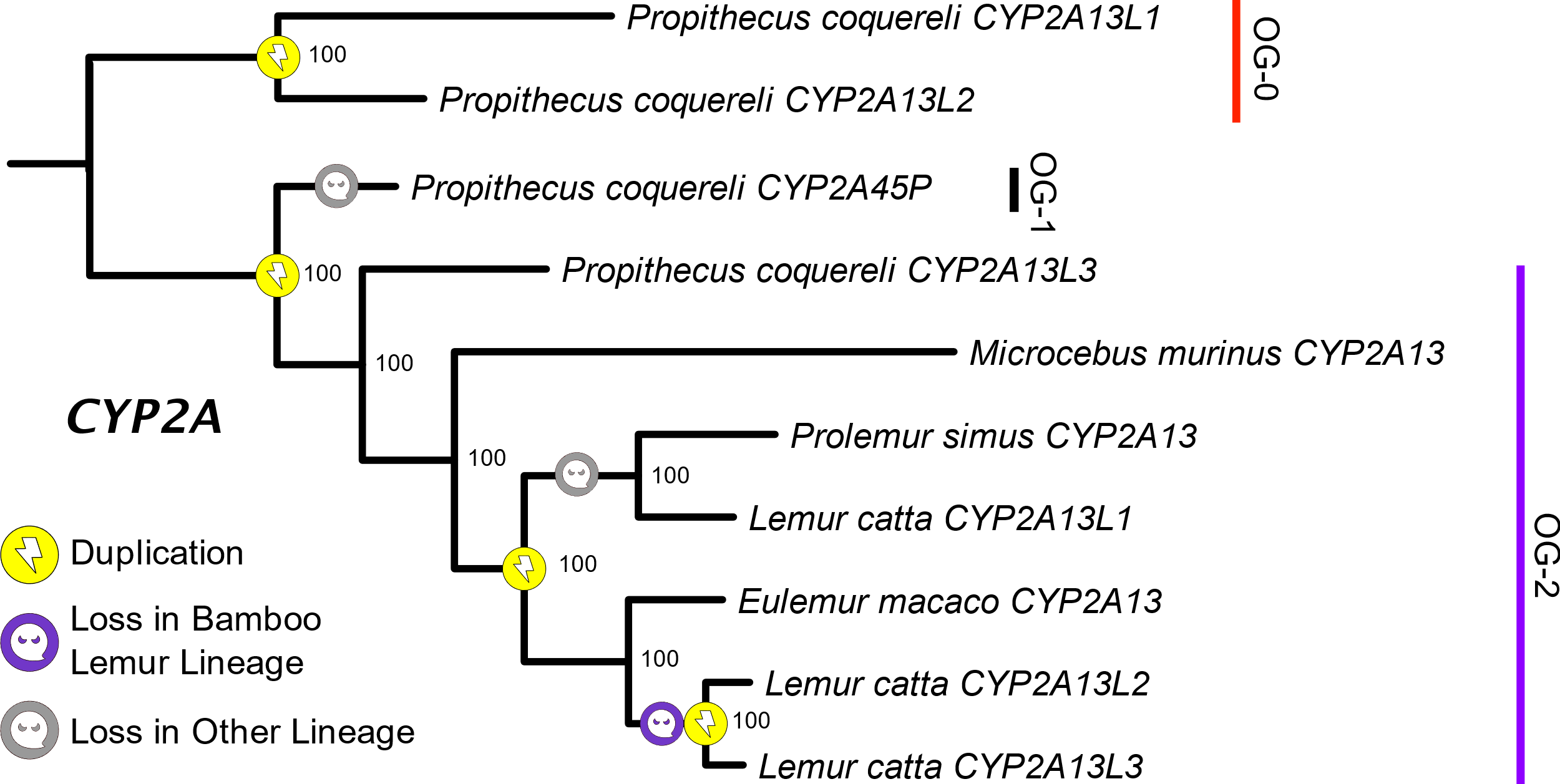

Figure S3: Phylogenetic tree for the *CYP2A* subfamily among five species of lemur.

Bootstrap values are displayed as percentages. Duplications and orthogroups in this tree were defined or confirmed by Possvm.

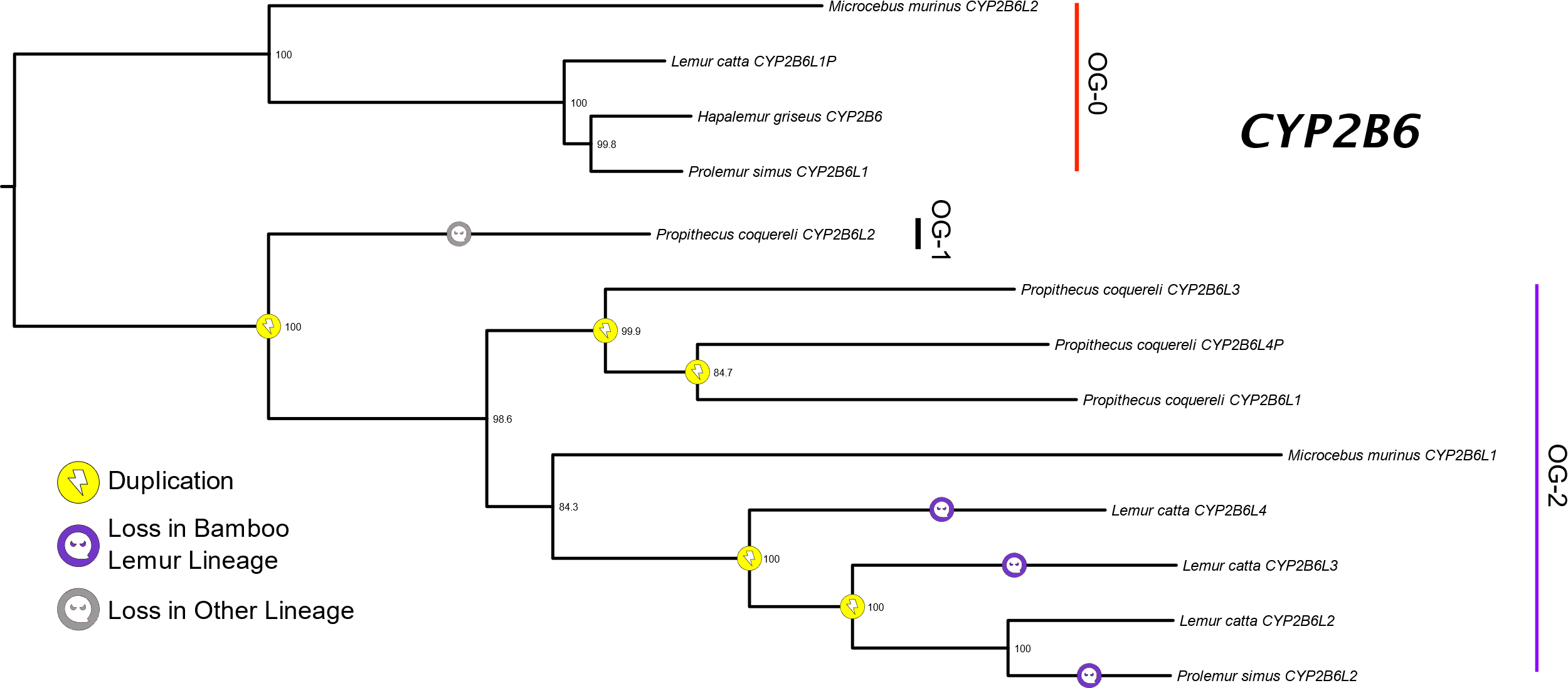

Figure S4: Phylogenetic tree of *CYP2B6* homologs among five species of lemur.

Duplications and orthogroups in this tree were defined or confirmed by Possvm. The software automatically midpoint-roots trees as part of its algorithm, which led to the position of *Prop. coquereli CYP2B6L2* as displayed here rather than an alternate interpretation, which would still be compatible with this tree, of this gene as part of the OG-0 clade. This scenario would be more parsimonious than what is displayed here because it would posit zero loss events along the branch leading to *Prop. coquereli CYP2B6L2*.

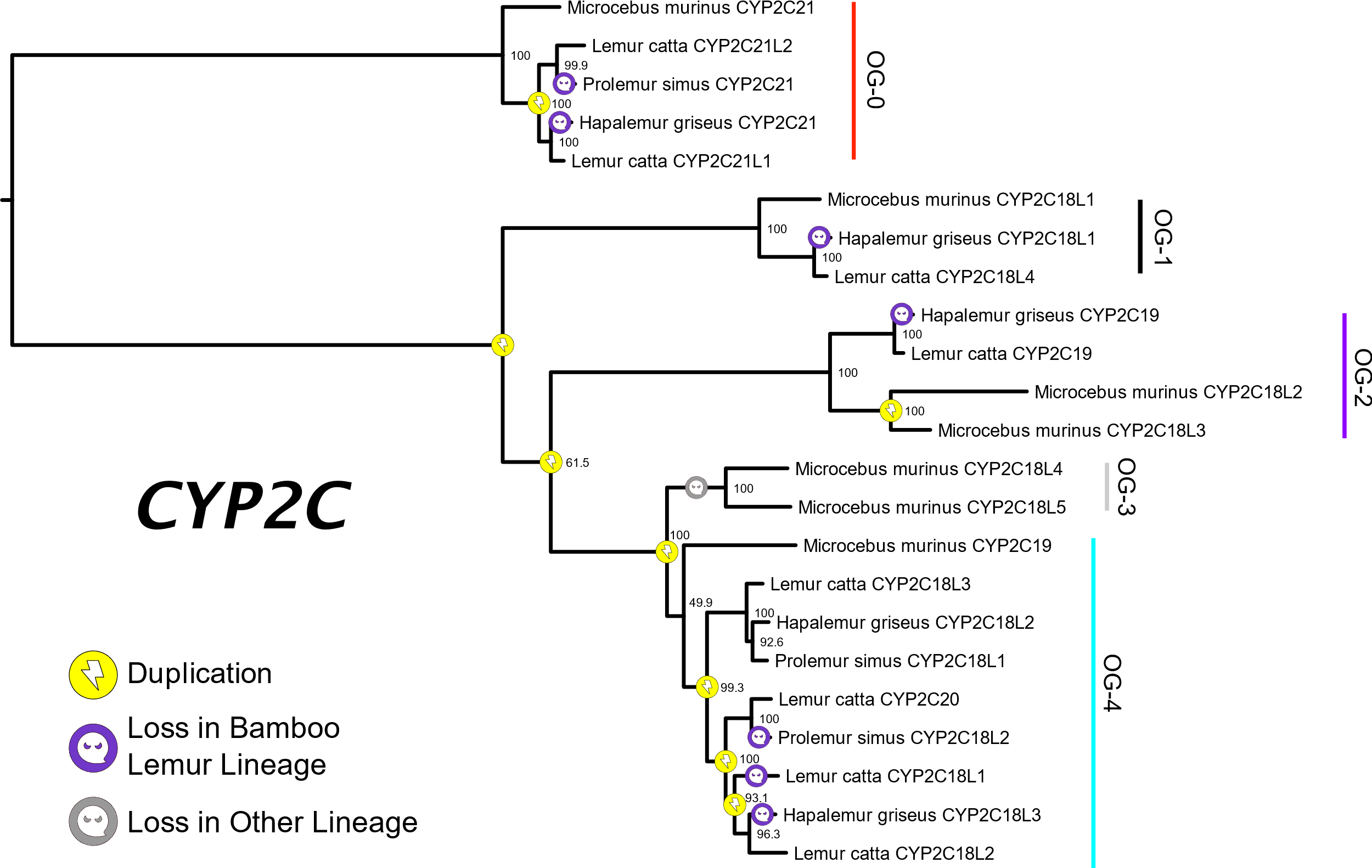

Figure S5: Phylogenetic tree of the *CYP2C* subfamily among four species of lemur.

Bootstrap values are displayed as percentages. Duplications and orthogroups in this tree were defined or confirmed by Possvm.

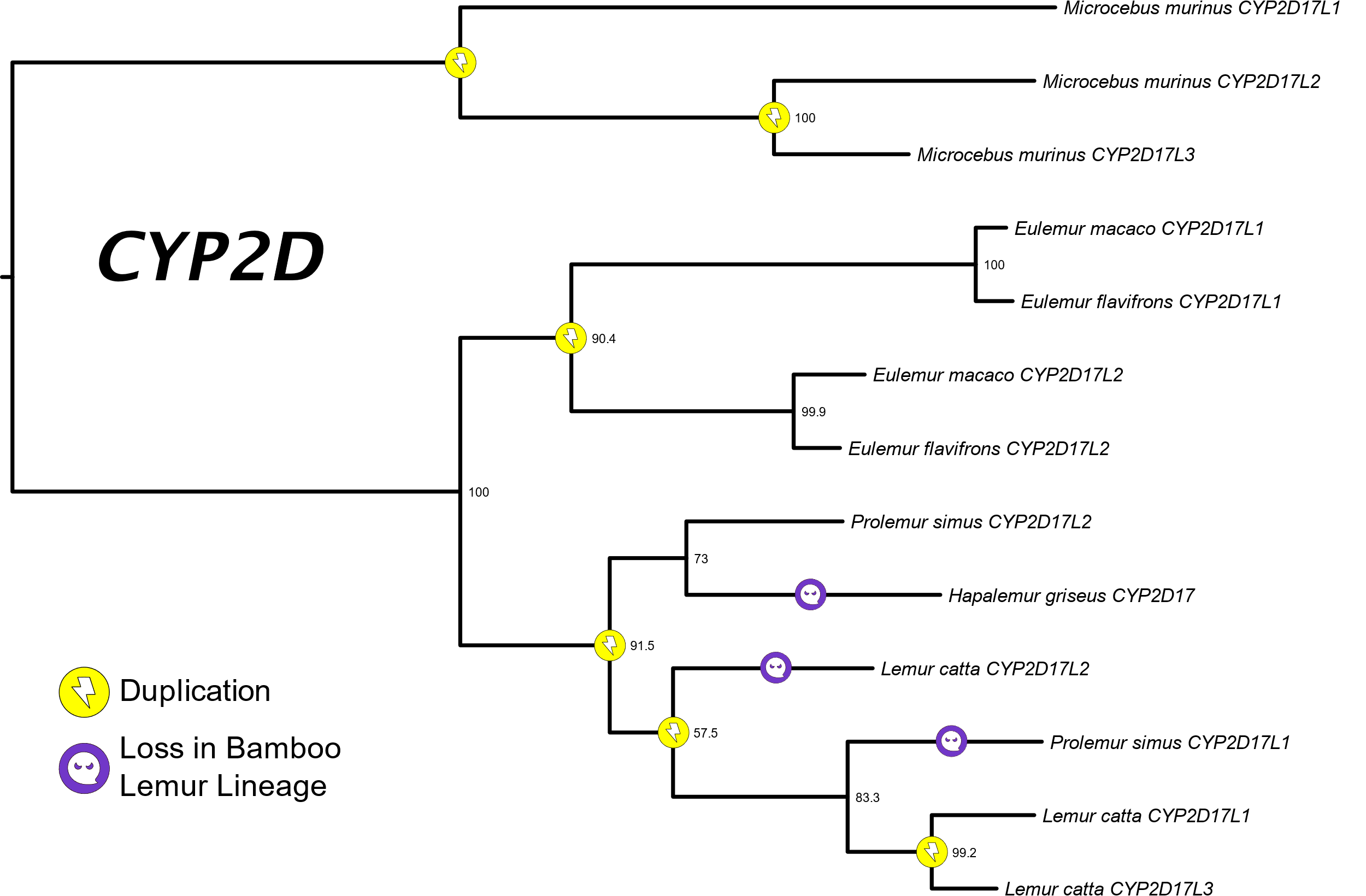

Figure S6: Phylogenetic tree of the *CYP2D* subfamily among six species of lemur.

The loss specified on the branch leading to *H. griseus CYP2D17* refers to a potential pseudogenization event of that gene itself because premature stop codons were detected in all three ORFs within some of the 3’ portion of this gene’s transcript. Bootstrap values are displayed as percentages. Duplications in this tree were defined by Possvm.

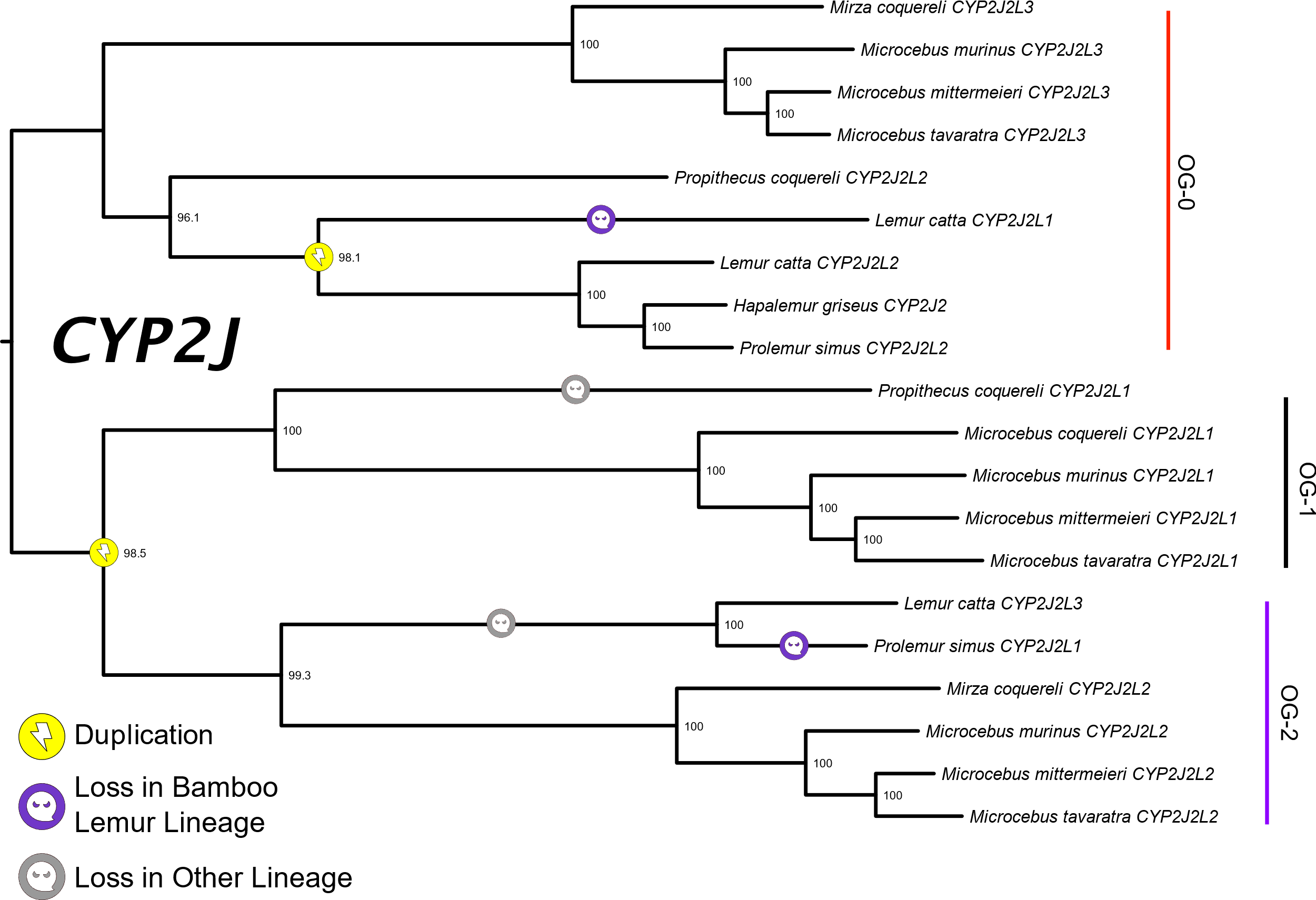

Figure S7: Phylogenetic tree of the *CYP2J* subfamily among seven species of lemur.

Bootstrap values are displayed as percentages. Duplications and orthogroups were detected or confirmed by Possvm.

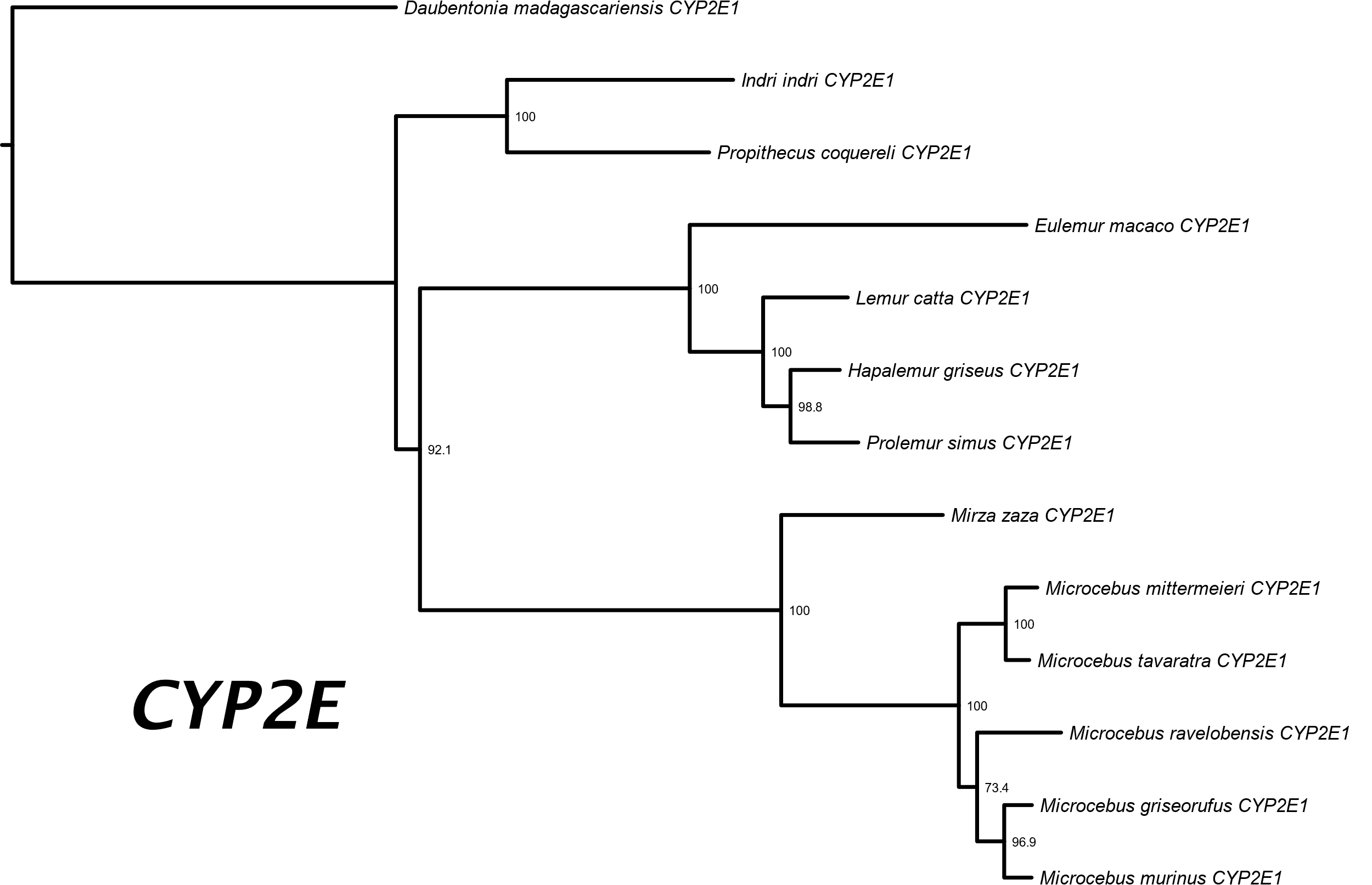

Figure S8: Phylogenetic tree for the *CYP2E* subfamily among 13 species of lemur.

Bootstrap values are displayed as percentages.

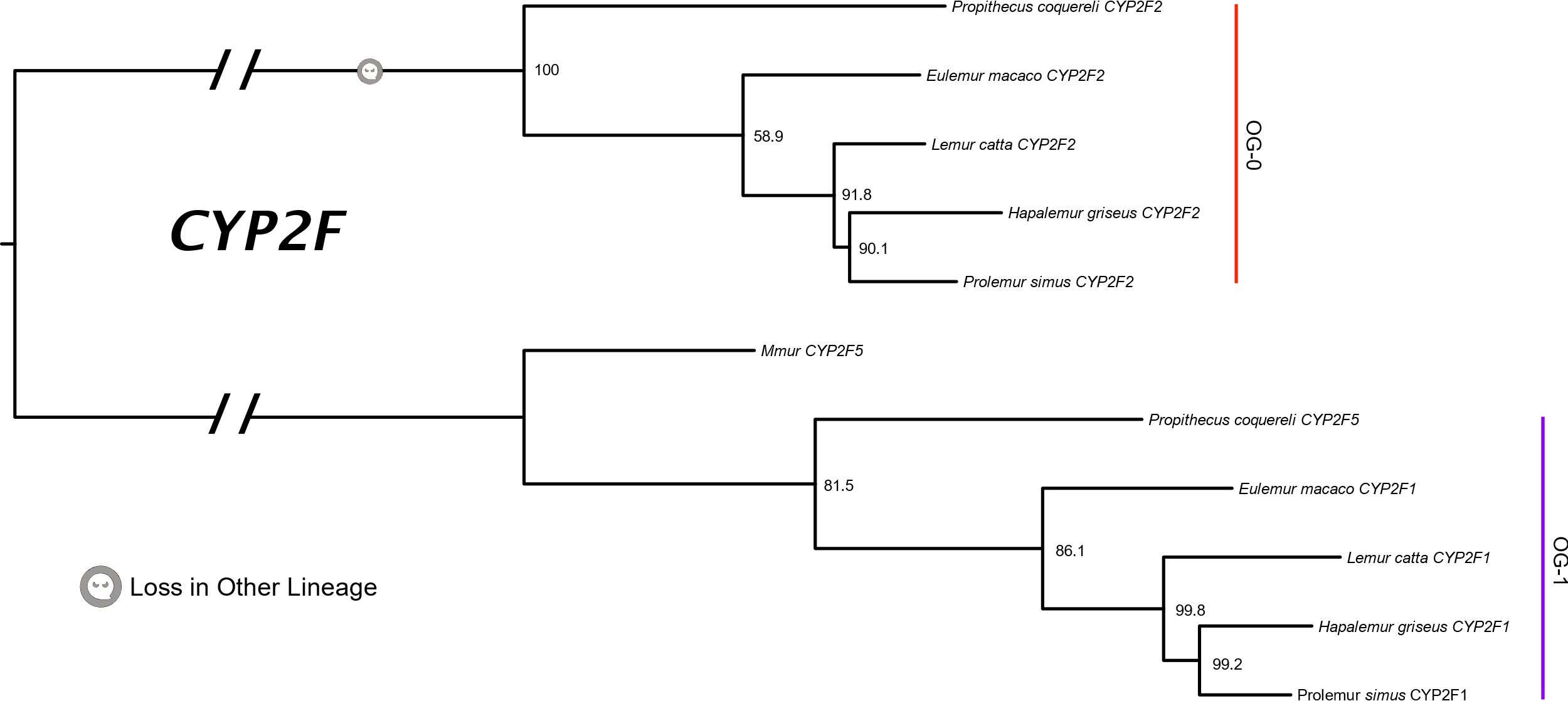

Figure S9: Phylogenetic tree for the *CYP2F* subfamily among six species of lemur.

The single deletion in this tree refers to the loss of a CYP2F2 ortholog for Mic. murinus. Bootstrap values are displayed as percentages. Orthogroups in this tree were defined or confirmed by Possvm.

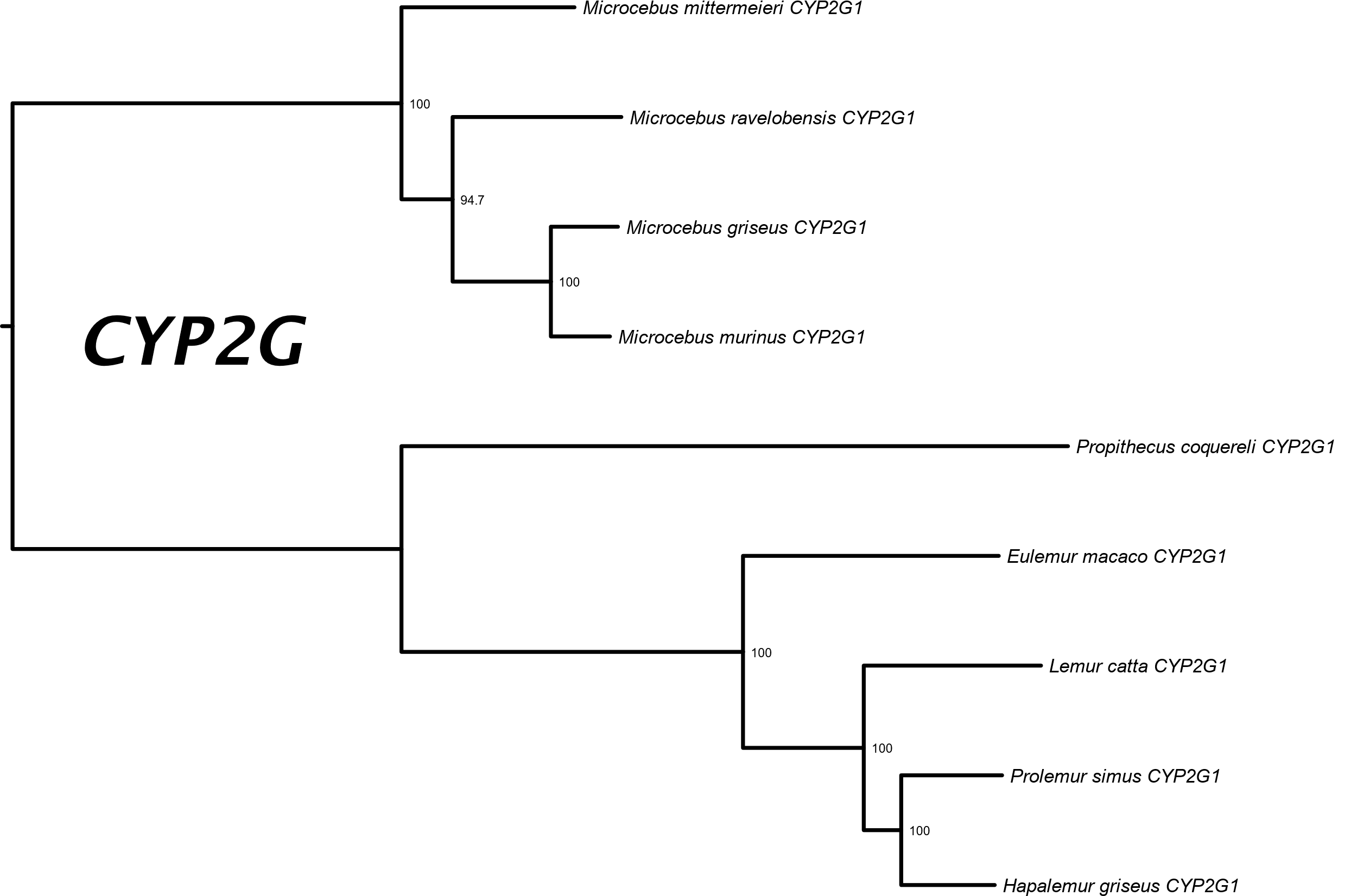

Figure S10: Phylogenetic tree for the *CYP2G* subfamily among nine species of lemur.

Bootstrap values are displayed as percentages.

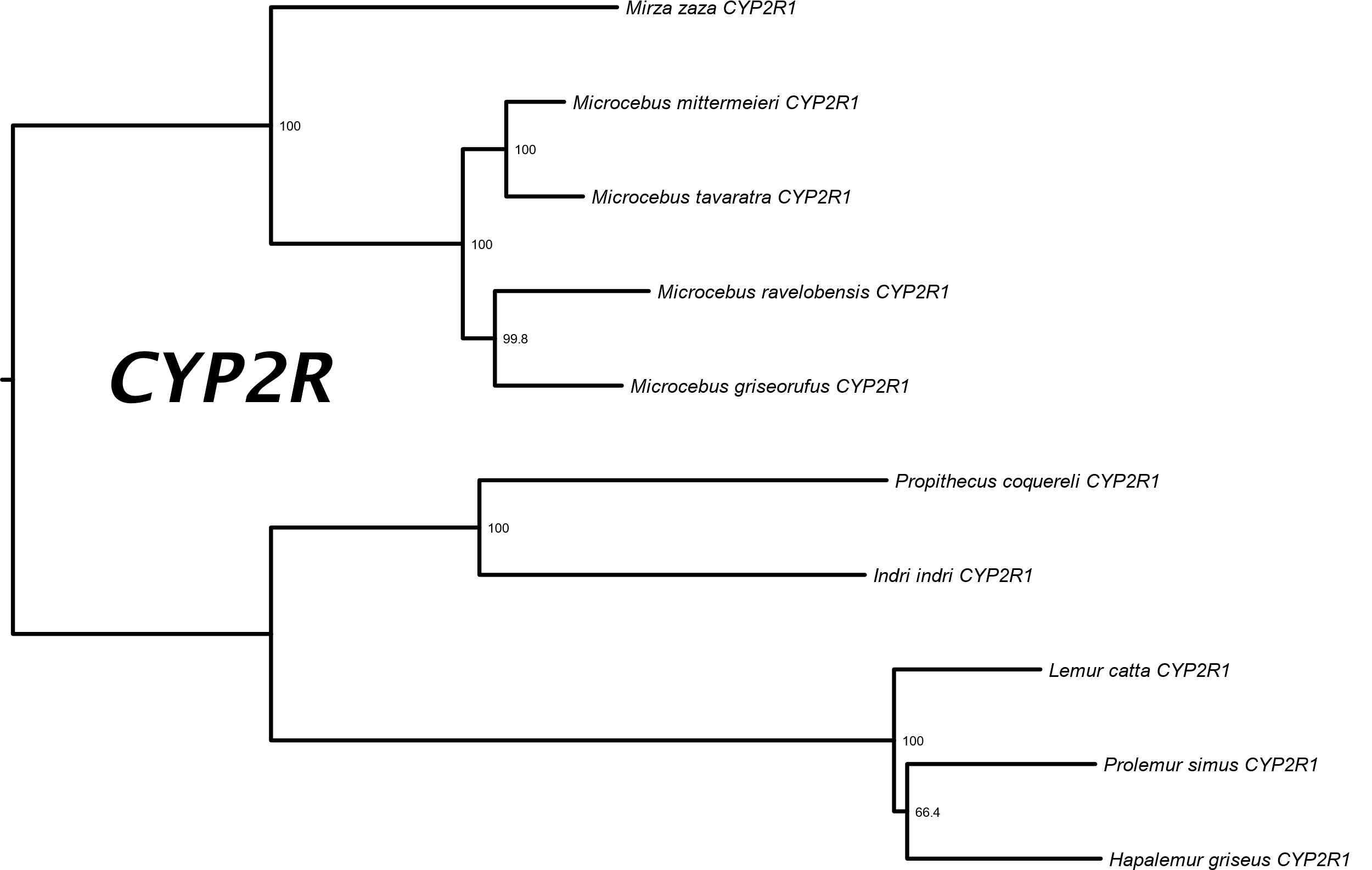

Figure S11: Phylogenetic tree for the *CYP2R* subfamily among 10 species of lemur.

Bootstrap values are displayed as percentages.

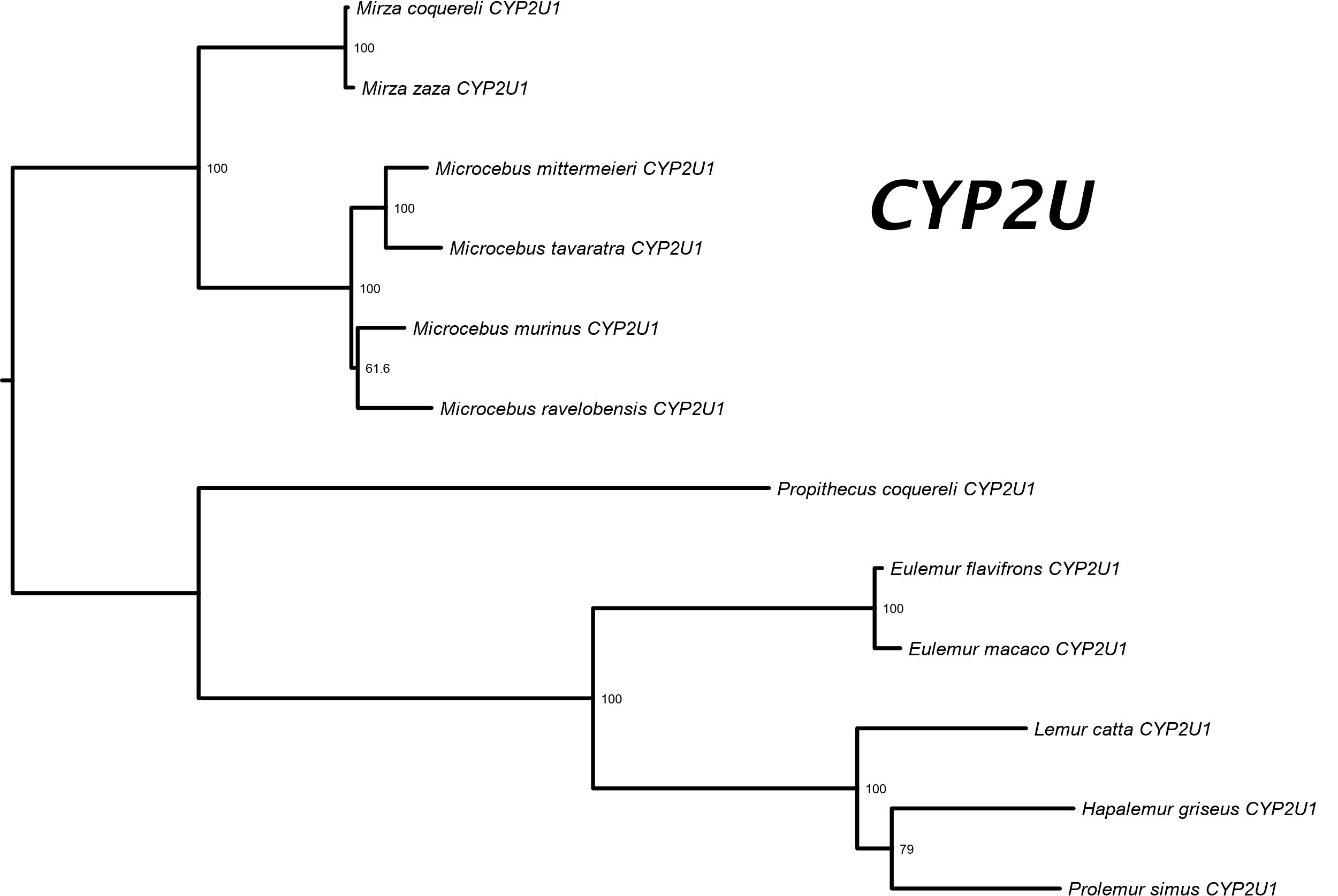

Figure S12: Phylogenetic tree for the *CYP2U* subfamily among 12 species of lemur.

Bootstrap values are displayed as percentages.

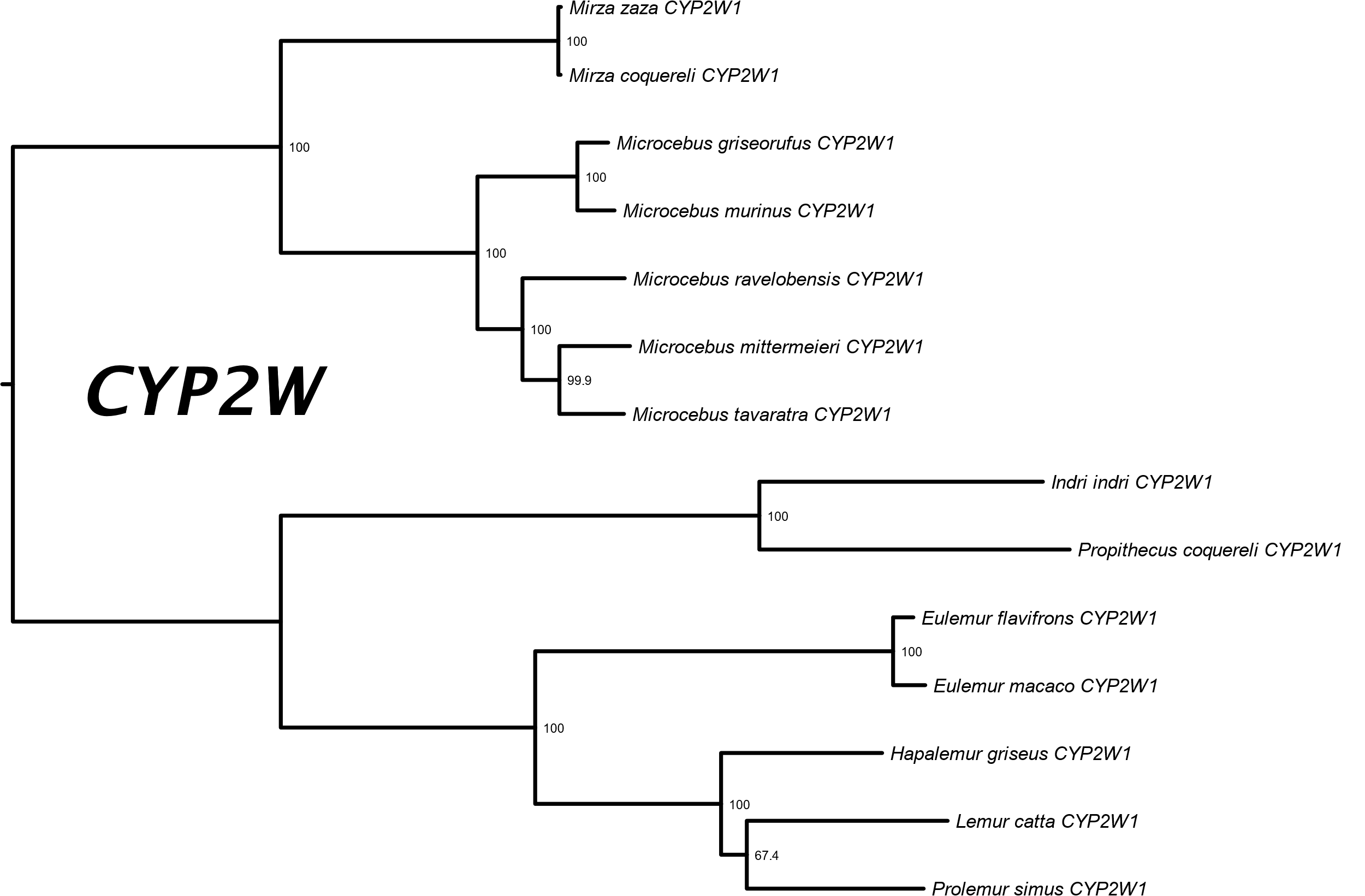

Figure S13: Phylogenetic tree of the *CYP2W* subfamily among 14 species of lemur.

Bootstrap values are displayed as percentages.

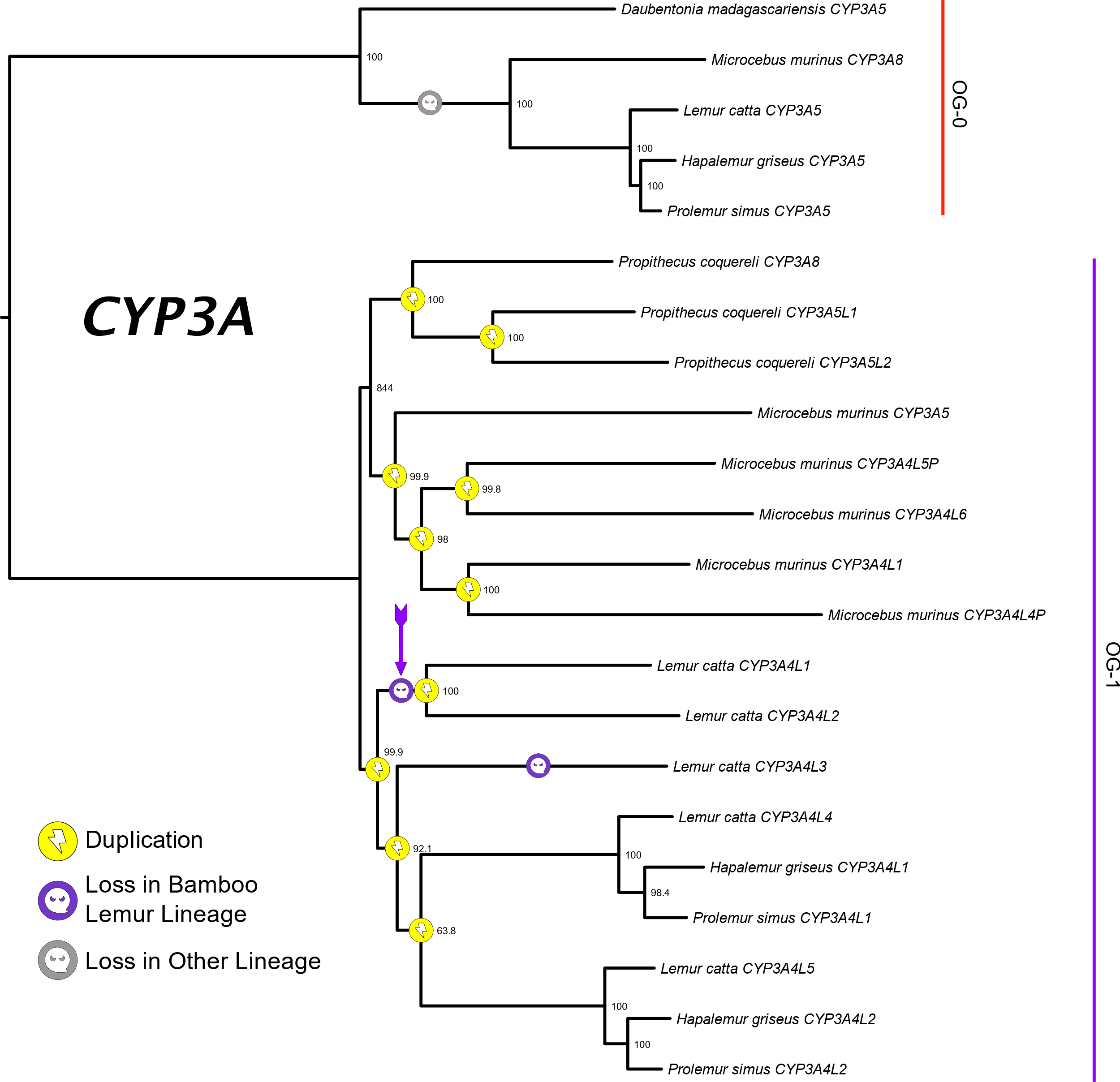

Figure S14: Phylogenetic tree of the *CYP3A* subfamily among six species of lemur.

The loss event indicated by the violet arrow is the most parsimonious interpretation for the number and timing of loss(es) in this part of OG-1 but a more complicated interpretation where two losses occurred in the terminal branches for *L. catta CYP3A4L1* and *CYP3A4L2* would not necessarily be falsified by this tree in light of those branches’ relatively large branch lengths. Bootstrap values are displayed as percentages.

Table S3: Results from hypothesis tests using RELAX.

| Gene Group | Likelihood Ratio | *p* | *K* |
| --- | --- | --- | --- |
| 1A | 0.61 | 0.433 | 1.17 |
| 1B | 0.86 | 0.355 | 0.42 |
| 2A | 2.56 | 0.109 | 0.61 |
| 2B | 0.40 | 0.527 | 0.64 |
| 2C | 29.1 | 6.8e-8* | 1.42† |
| 2D | 1.67 | 0.196 | 5.12 |
| 2E | 0.13 | 0.716 | 0.90 |
| 2F | 26.3 | 3.0e-7* | 0.00 |
| 2G | 0.05 | 0.819 | 1.12 |
| 2J | 5.70 | 0.017* | 0.51 |
| 2R | 0.79 | 0.376 | 0.65 |
| 2S | 1.05 | 0.306 | 0.65 |
| 2U | 0.49 | 0.483 | 1.46 |
| 2W | 0.03 | 0.871 | 1.04 |
| 3A | 2.91 | 0.088 | 0.55 |

*: *p* < 0.05

†: Follow-up analysis by aBSREL () and BUSTED (), both run with synonymous-rate variation (), showed that this intensification signal was driven by a site in the *CYP2C21* gene from our *Hapalemur griseus* assembly (*i.e.,* Hgris_2C21, see below). This signal was not shared by any of the other *Hapalemur* orthologs for this gene.

Figure S15: Region of the *H. griseus CYP2C21* ortholog responsible for the significantly heightened *k* estimate for *CYP2C*.

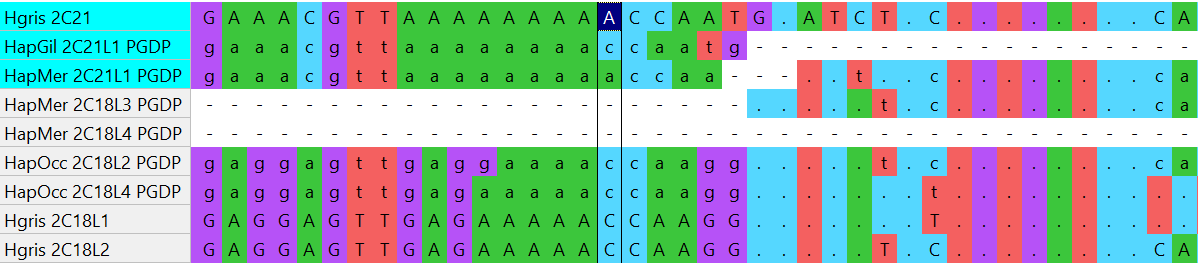

Table S4: Differences of nonsynonymous and synonymous ratios (*dN –* *­dS*) for *CYP2F1.*

|  | 1 | 2 | 3 |
| --- | --- | --- | --- |
| 1. *Prolemur-Lemur* Ancestral Sequence |  |  |  |
| 2. *Prolemur-Hapalemur* Ancestral Sequence | -0.000939 |  |  |
| 3. Hgris_2F1 | -0.001999 | -0.001071 |  |
| 4. Psim_2F1 | -0.009424 | -0.008363 | -0.010982 |

Table S5: Differences of nonsynonymous and synonymous ratios (*dN –* *­dS*) for *CYP2F2.*

|  | 1 | 2 | 3 | 4 | 5 |
| --- | --- | --- | --- | --- | --- |
| 1. *Prolemur-Lemur* Ancestral Sequence |  |  |  |  |  |
| 2. *Prolemur-Hapalemur* Ancestral Sequence | 0.000914 |  |  |  |  |
| 3. Psim_2F2 | -0.006614 | -0.007538 |  |  |  |
| 4. *H. griseus-gilberti* Ancestral Sequence | 0.016051 | 0.015097 | 0.007597 |  |  |
| 5. HapGil_2F2_PGDP | 0.018335 | 0.017098 | 0.013807 | 0.002420 |  |
| 6. Hgris_2F2 | 0.015449 | 0.014486 | 0.006845 | 0.000182 | 0.002556 |
